# Pcdh18a-positive tip cells instruct notochord formation in zebrafish

**DOI:** 10.1101/257717

**Authors:** Bernadett Bosze, Benjamin Mattes, Claude Sinner, Kathrin Stricker, Victor Gourain, Thomas Thumberger, Sham Tlili, Sabrina Weber, Joachim Wittbrodt, Timothy E. Saunders, Uwe Strähle, Alexander Schug, Steffen Scholpp

## Abstract

The notochord defines the axial structure of all vertebrates during development. Notogenesis is a result of major cell reorganization in the mesoderm, the convergence and the extension of the axial cells. However, it is currently not known how these processes act together in a coordinated way during notochord formation. Analysing the tissue flow, we determined the displacement of the axial mesoderm and identified, relative to the ectoderm, an actively-migrating notochord tip cell population and a group of trailing notochordal plate cells. Molecularly, these tip cells express Protocadherin18a, a member of the cadherin superfamily. We show that Pcdh18a-mediated recycling of E-cadherin adhesion complexes transforms these tip cells into a cohesive and fast migrating cell group. In turn, these tip cells subsequently instruct the trailing mesoderm. We simulated cell migration during early mesoderm formation using a lattice-based mathematical framework, and predicted that the requirement for an anterior, local motile cell cluster could guide the intercalation of the posterior, axial cells. Indeed, grafting experiments validated the predictions and induced ectopic notochord-like rods. Our findings indicate that the tip cells influence the trailing mesodermal cell sheet by inducing the formation of the notochord.

## Introduction

The notochord is the most prominent hallmark of the phylum Chordata and serves as the common embryonic midline structure for all their members, including humans. Serving as an embryonic scaffold for the surrounding mesoderm to subsequently form the skull, the membranes of the brain, and, most importantly, the vertebral column [1], it plays a central role in the genesis of the vertebral body. Generation of the dorsal mesoderm, including the notochord, is based on immense cellular rearrangements and the zebrafish has been proven to be an excellent *in vivo* system to study this process. During embryogenesis, the notochord originates from the dorsally localized shield organizer of the embryos when cells break their cell-cell junctions and undergo single-cell ingression at the onset of gastrulation in zebrafish [2]. After internalization, these mesodermal progenitor cells organize into a coherent cell sheet to migrate from the embryonic margin towards the animal pole of the embryo. This cell sheet can be subdivided into the anterior prechordal plate mesoderm (PPM), the axial notochordal plate, and the lateral plate mesoderm (LPM) [3]. Subsequently, the notochordal plate narrows and elongates in the perpendicular axis to finally transform into the notochord, whereas the LPM cells migrate towards the midline to form the remaining mesodermal organs. The molecular mechanisms controlling these cell movements, generally known as convergence and extension, have been extensively studied in the LPM [4,5]. Cadherin-based junctions mediate collective cell polarization and generate the common force that organizes the mesodermal plate in vertebrates [6]. Wnt-planar cell polarity (PCP) signalling controls the internalization of E-cadherin cell adhesion complexes to coordinate cell movements [7]. Directed migration is a prerequisite for coordinated cohort movement, and a polarized distribution of the Wnt-PCP component Vangl2 has been recently observed in the zebrafish LPM [8]. Therefore, a hypothesis for notochord formation suggests that, similar to the LPM, cell migration leads to the intercalation and elongation of the notochord plate cells to form the rod-shaped notochord [9,10]. An alternative hypothesis highlights the importance of the LPM and suggests that the notochordal plate intercalates as a consequence of LPM pushing forces [9].

Here, we identified a cell group located at the tip of the zebrafish notochord, which shows enhanced endocytosis dynamics of E-cadherin regulated by prominent Pcdh18a expression. Pcdh18a/E-cadherin-positive notochord tip cells show unique properties allowing them to cluster and migrate faster thus affecting the transformation of the trailing axial mesoderm into the notochord.

## Results

### Tip cells are required to shape the zebrafish notochord

We mapped the expression of key cell adhesion molecules relative to the expression of mesodermal marker genes to characterize the axial mesoderm during early notochord development in zebrafish. We identified a cell cluster that expresses *pcdh18a, e-cad*, and *fzd7a* (Figure 1A; supplementary figure 1A-G). This cell cluster contains approximately 70-80 cells and is localized in the axial mesoderm between the e-cad/gsc-positive PPM and the *ntl/gsc-positive* notochord (Figure 1B, Supplementary figure 1C). Due to their prominent position, we refer to this confined cell population as the notochord tip cells (NTC) in the remainder of the text. Nodal signalling is required for the induction and involution of the mesoderm-derived notochord at the shield organizer [10,11]. After blocking the Nodal-signalling cascade, we observed a strong down-regulation of *pcdh18a* expression, suggesting that Nodal-mediated organizer signalling controls *pcdh18a* expression in the NTC (Figure 1C). Pcdh18a is a transmembrane protein that displays dynamic endocytic cycling between the cell membrane and vesicles (Figure 1D).

**Figure 1:**
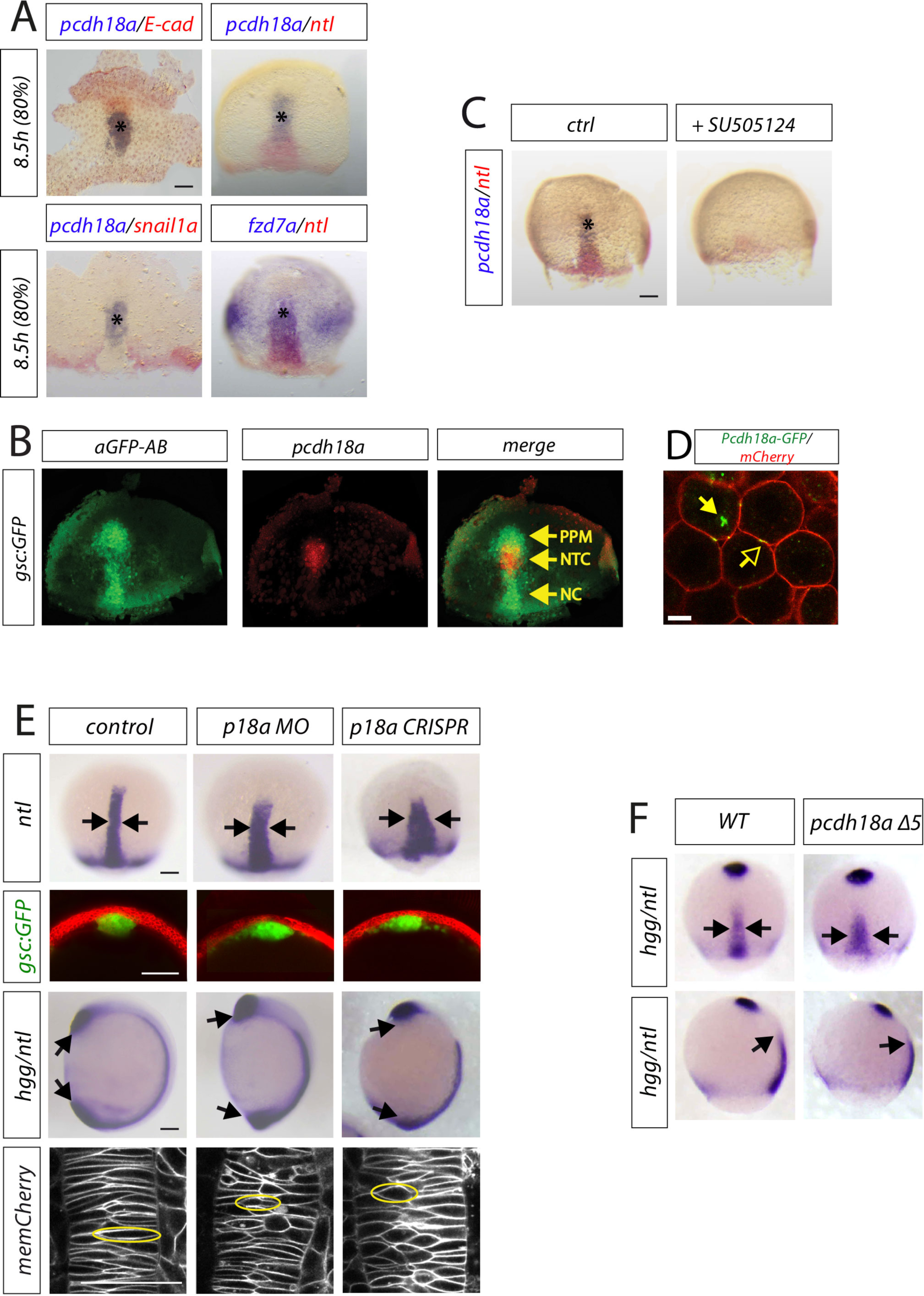
A cell group in the axial mesoderm regulates notochord formation in zebrafish. A, Transcriptional characterization of the notochord tip cells (NTC) by whole-mount double in situ hybridization (ISH) of WT zebrafish embryos with the indicated markers. B, Mapping of pcdh18a expression relative to GFP expression in tg(gsc:GFP). Notably, gsc:GFP labels the prechordal plate mesoderm (PPM) and the trailing notochord (NC). C, Inhibition of Nodal signaling by SU505124 treatment (30 μM) from 4 - 8.5 hpf. D, Confocal imaging of living zebrafish embryo at 5 hpf showing subcellular localization of Pcdh18a-GFP. GPI-mCherry marks cell membranes. Arrows indicate punctae of Pcdh18a localization at the membrane (open arrows) and intracellularly (closed arrows). Scale bar 10μm. E, WT embryos or tg(gsc:GFP) embryos were injected with pcdh18a-MO (0.5 mM) or with preincubated pcdh18a-gRNA (0.2 ng) & codon-optimized Cas9-mRNA (0.1 ng). Morpholino-based knockdown of pcdh18a (MO) or CRISPR-based knockout in zebrafish leads to a wider and shorter notochord marked by ntl expression at 9 hpf (arrows). See Supplementary figure 2 for control experiments. Analysis of the shape of the notochord in a cross-section of gsc:GFP transgenic embryos that were injected with the indicated constructs at 10 hpf. Scale bar, 100 μm. At 11 hpf, the body length was significantly shorter in the Pcdh18a-deficient embryos, as shown in an ISH-based analysis of notochord hgg/ntl (MO: 62%, CRISPR: 84%; arrows). Confocal microscopy-based analysis of cell shapes in the notochord of embryos that were microinjected with the indicated constructs at 12 hpf (cells with exemplary morphology were surrounded with a yellow circle). See Supplementary figure 3G for quantification. F, *pcdh18a A5* mutant embryos show a wider and shorter axial mesoderm (width 50.8μm / length 141μm, n=4) compared to wt / heterozygous mutant embryos (width 41.5μm / length 198.4μm, n=4) of the same clutch. See Supplementary figure 2 for details.

To analyse whether Pcdh18a is required for mesoderm organization in the gastrula, we altered the Pcdh18a levels using a Morpholino (MO)-based antisense approach [12] and we complemented this analysis with a CRISPR/Cas9-based knockout strategy [13] (Figure 1E; supplementary figure 2B-F). We found that a reduction of *pcdh18a* expression led to a wider axial mesoderm, as shown by the lateral expansion of the *ntl* expression domain and of the *gsc* domain in *tg(gsc:GFP)* zebrafish embryos (Figure 1E). Furthermore, we observed that blockade of Pcdh18a function led to a significant reduction in the extension of the axial mesoderm, similar to embryos that ectopically express E-cad and p120 (Supplementary figure 3F). In parallel, we generated a genetic mutant using the CRISPR/Cas9 system. We detected a 5bp deletion in exon1 of the Pcdh18a gene (Supplementary Fig. 2G). These *pcdh18a* mutants exhibit a wider and shorter notochord shown by the expression domain of *ntl* and *hgg* at 9hpf (Figure 1F). *pcdh18a* mutant, crispant and morphant embryos, therefore, show a similar phenotype. At 26 hpf, the head and trunk display a WT phenotype; however, the tail shows a strong deformation, suggesting a compromised axial scaffold (Supplementary figure 3F). Remarkably, Pcdh18a function in notochord formation is largely independent of the convergent extension of the LPM (Supplementary figure 4B,C).

**Figure 2:**
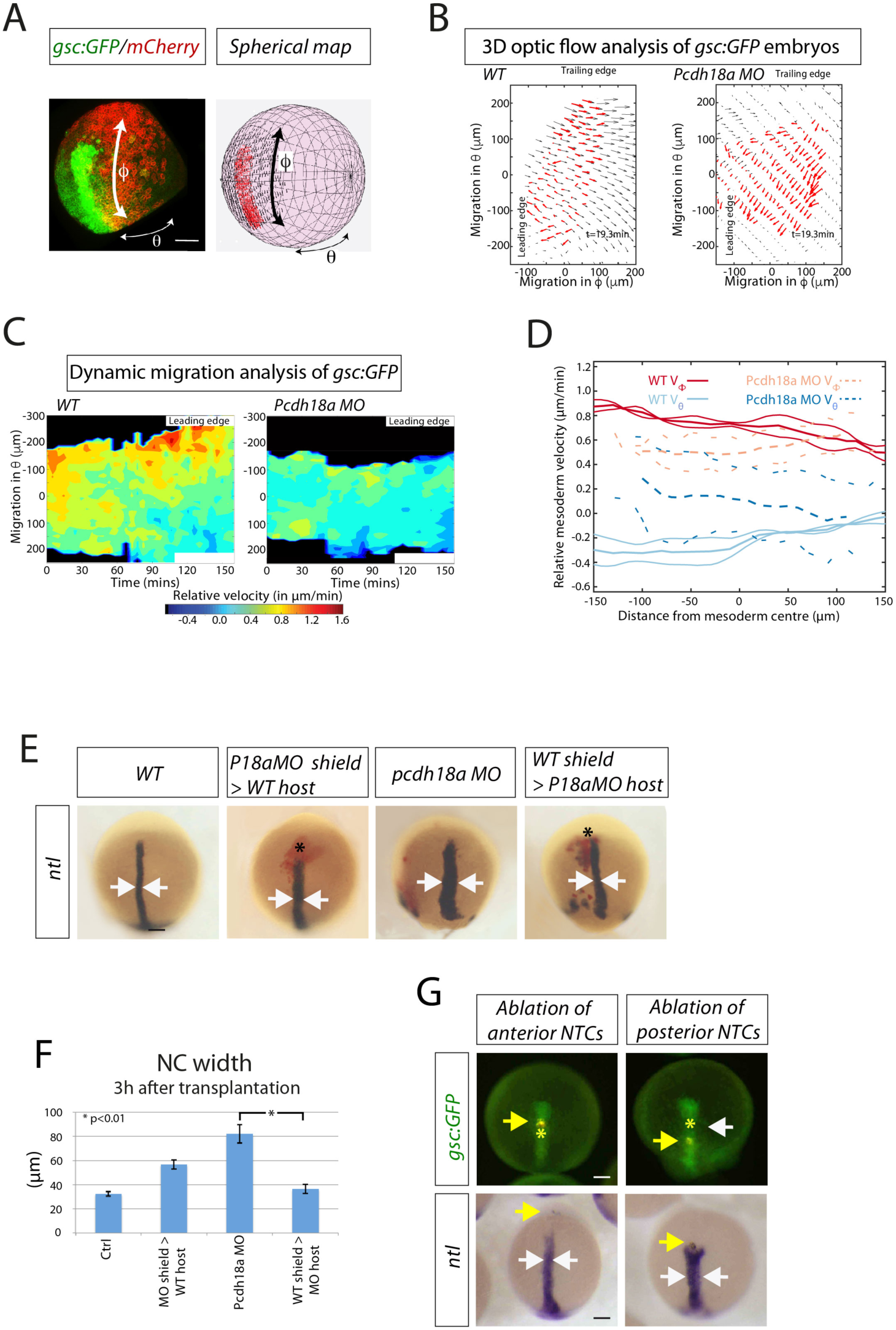
Analysis of the influence of the NTC on notochord formation. A-D, 3D-optic flow analysis. A spherical mapping of the optic flow of the fluorescent signal of a *gsc:GFP/memCherry* embryo onto a spherical coordinate system with θ as azimuth angle and Φ as polar angle. B, 2D Mercator projection of flow field of mesoderm (red arrows) compared with ectoderm above (black arrows). Both embryos analysed at 5 hpf. C, The corresponding heat map kymograph shows the relative velocity of the mesoderm roughly every 5min. D, Thick solid (dashed) lines are smoothed average profiles of WT (Pcdh18a deficient) embryos, with individual embryo profiles shown in lighter lines in Φ (red/orange) and θ (light/dark blue) directions. E, pcdh18a MO donor shield was transplanted into WT hosts and vice versa. After 3h, the embryos were fixed and subjected to ISH against *ntl.* Donor cells are marked in red. Arrows indicate the width of the notochord. F, Quantifications display mean value, standard error of mean (SEM), and significance level of six independent embryos per experiment as indicated. G, Ablation of cell rows anterior (5th GFP positive cell row) or posterior (15th GFP positive cell row) to the NTC in the *tg(gsc:GFP)* fish line. Embryos were injected with a nuclear marker (Histone 2B-mCherry) and cell rows were ablated at 7 hpf using ultrashort laser pulses of a two-photon microscope. Embryos were raised to 10 hpf, fixed, and subjected to ISH against ntl. After ablation of the anterior NTC row, embryos develop an elongated notochord (n=11/11), whereas the notochord progenitor cells move slower and a gap appears towards the anterior NTC in embryos with ablation of a cell row in the posterior NTC (white arrow). Consequently, the trailing ntl expression domain remains shorter and broader (n=6/10). Yellow arrows mark the ablated cell rows. Asterisks mark the location of pcdh18a-positive NTC. Scale bar, 100 µm.

During notogenesis, central mesodermal cells take on a bipolar shape and intercalate mediolaterally and pull together (convergence) [14]. In zebrafish, polarized radial intercalations of axial mesoderm separates anterior and posterior neighbours, and lengthening the field along the AP axis (extension) [9,10]. Therefore, we investigated the shape of the notochord cells in Pcdh18a deficient embryos at 12hpf (6 somite-stage). We found that the cells had a less bipolar shape and displayed a more circular form in embryos with reduced Pcdh18a levels (Figure 1E; Supplementary figure 3G), suggesting that mediolateral narrowing and AP lengthening of axial mesoderm is reduced. Next, we investigated the migratory properties of the central mesoderm. We performed a three-dimensional optic flow analysis; 3D-KLT, [15] in zebrafish embryos to obtain local tissue velocity measurements of both the mesoderm and ectoderm, see Methods for details. To aid quantification, we mapped the flow of the migrating tissues onto a sphere (obtained from a spherical fit of the embryo shape) and calculated velocities in spherical coordinates. The sphere’s equator and poles were oriented such that the main axis of elongation of the mesoderm was aligned with the meridian (Figure 2A). For the flow analysis of the forming notochord, we used time-lapse movies from *gsc:GFP* embryos injected with memCherry and scanned from 5hpf - 8hpf (Figure 2A,B Supplementary figure 3A). In this way, we calculated the velocity field for the *gsc:GFP* positive axial mesoderm and the ectoderm above in WT and Pcdh18a morphant embryos (Figure 2B). From these, we generated kymographs of the relative velocity between the mesoderm and ectoderm (Figure 2C, Supplementary figure 3B). We found that in WT embryos the leading edge and trailing edge of the mesodermal layer have distinct dynamics with respect to the underlying ectoderm: the leading cell population migrates faster (red-yellow areas) towards the animal pole compared to the trailing notochordal plate (green-blue areas) in the φ direction (Figure 2C). The drop in relative migration speed from animal to vegetal becomes more apparent over time (after 150min). Next, we compared the mesoderm dynamics in WT embryos to Pcdh18a deficient embryos. We found that the tip cells migrate slower in the φ direction in Pcdh18a deficient embryos (Fig. 2C,D, Supplementary figure 3C). In contrast, we found that migration in the θ direction is enhanced in the tip cells of morphants (Figure 2D), whereas it is reduced in the trailing edge (Supplementary Figure 3D). We hypothesized that pronounced elongation (φ migration) mediated by Pcdh18a function in the NTCs contributes to enhanced convergence (θ migration) of the follower cells - the notochordal plate. To test whether Pcdh18a-positive tip cells influence the organization of the trailing notochord, we altered the Pcdh18a levels in the NTC in a cell transplantation experiment (Figure 2E,F). We discovered that the notochord of host embryos carrying Pcdh18a-deficient NTC is wider than the control embryos. In a reverse experiment, we introduced the NTC of WT origin into Pcdh18a-deficient embryos. The axis of *pcdh18a* morphant embryos with WT NTC was significantly thinner compared to the *pcdh18a* morphant embryos. Based on the fine mapping of the NTC in the *tg(gsc:GFP)* embryos (Supplementary figure 4A), we performed a pulsed laser-ablation experiment in which we ablated a cell row between the PPM and the NTC (arrow 1) or between the NTC and the notochord (arrow 2). Ablation of the cells in the anterior PPM did not lead to an obvious alteration in the morphology of the chordamesoderm (Figure 2G). However, ablation of a cell row in the posterior NTC led to the formation of a gap between the NTC and the notochordal plate (white arrow), and these embryos displayed a shorter and wider notochord. These data suggest that the NTC guide the posterior notochord. In summary, these data suggest that the Pcdh18a-positive NTC constitute a border cell cluster which organizes the elongation and intercalation of the posteriorly trailing notochord.

### Pcdh18a regulates E-cad/Fzd7a endocytic recycling

Cadherins accumulate at cell-cell contact sites, such as adherens junctions, to regulate cell migration and cell adhesion [16], similar to protocadherins, which influence homophilic and heterophilic interactions of cadherins [17,18]. We therefore sought to determine a possible interaction between three identified NTC transmembrane proteins: Pcdh18a, E-cad, and Fzd7a. Co-localization and co-migration of Pcdh18a-mCherry with E-cad-GFP and Fzd7a-CFP was observed in zebrafish gastrula and in mouse fibroblasts that stably express E-cad-GFP (E-cad-GFP+ L cells) (Figure 3A; supplementary figure 5A-D).

**Figure 3:**
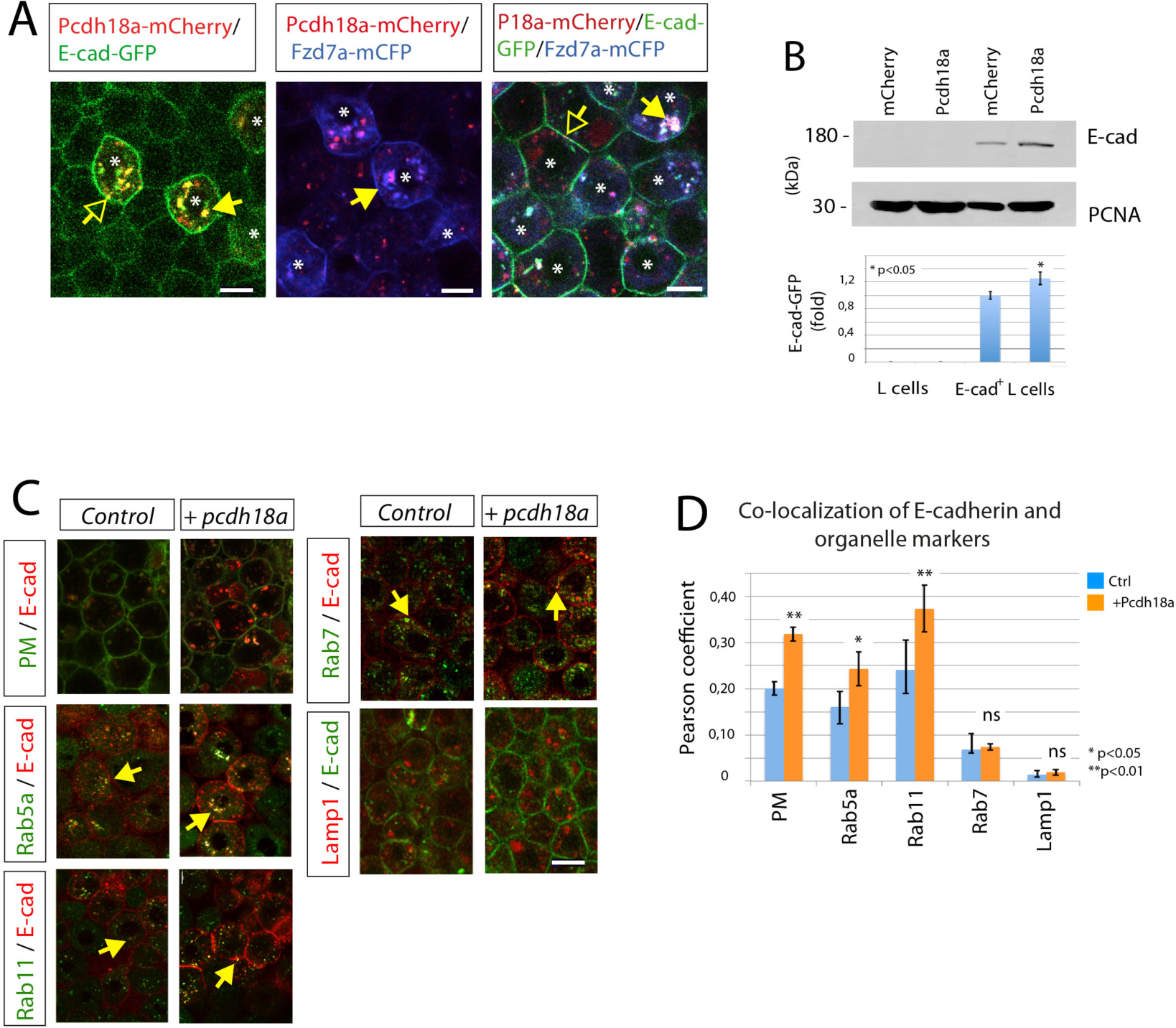
Pcdh18a regulates recycling of the E-cad/Fzd7a complex. A, Confocal images of zebrafish embryos at 5 hpf. Embryos were microinjected with 0.1 ng of mRNA for the indicated constructs and were imaged in vivo at 5 hpf. Pcdh18a is localized in the cell membrane (open arrows) and in endocytic vesicles (closed arrows), together with E- cad and Fzd7a. B, Quantification of the E-cad levels in the Pcdh18a-transfected L-cells. Equivalent amounts of lysates from murine L cells or stably E-cad-GFP-transfected L-cells that had been transfected with Pcdh18a were Western blotted and probed with an anti-GFP antibody; the results showed a 26% increase in the E-cad-GFP protein levels after Pcdh18a transfection. PCNA was used as a loading control. The experiments were performed in independent triplicate. C, Endocytic routing of E-cadherin at 50% epiboly. WT embryos and tg(rab5-GFP), tg(rab7-GFP), and tg(rab11-GFP) stable transgenic embryos were microinjected with 0.1 ng of the mRNAs for the indicated constructs. Arrows indicate E-cad localization with Rab proteins and Lamp1-positive vesicles. D, Pearson’s co-localization coefficient was calculated from 70 µm thick confocal stacks of 5 different embryos, each from (c). The error bars represent the SEM and significance, as indicated.

Remarkably, we observed increased E-cad-GFP fluorescence in vesicles and in the membrane in zebrafish blastula cells that co-express Pcdh18a (Figure 3A, supplementary figure 5A-C). A Western blot analysis of stably transfected E-cad-GFP+ L-cells also revealed an increase in E-cad expression in cells that co-express Pcdh18a (Figure 3B, Supplementary figure 5D). To determine which Pcdh18a-dependent mechanism might stabilize E-cad levels, as Pcdh18a function is dispensable for E-cad expression (Supplementary figure 9B), we tested whether Pcdh18a affects the subcellular routing of E-cad (Figure 3C). Using Pearson’s correlation coefficient, we detected increased localization of E-cad in the plasma membrane, in Rab5a early endosomes, and in Rab11 recycling endosomes when Pcdh18a was coexpressed (Figure 3D). However, co-localization changes with Rab7-positive late endosomes and Lamp1-positive lysosomes was below detection level. This suggests that Pcdh18a more effectively guides E-cad to the recycling pathway, hence enhancing the E-cad protein levels.

To support the hypothesis that Pcdh18a increases endocytic recycling of E-cad, an *in vivo* fluorescent recovery after bleaching (FRAP) of membrane-located E-cad-GFP was performed in zebrafish embryos at 5hpf. We found that the FRAP into a bleached region of E- cad-GFP expressing membrane is reduced in a WT embryo compared to that in an embryo co-expressing E-cad-GFP and Pcdh18a (Figure 4A). These data suggest that Pcdh18a facilitates recovery of fluorescence by E-cad-GFP by enhanced endocytic recycling to the membrane. Furthermore, we cannot exclude that Pcdh18a and Pcdh18a-ECD also influence motility of E-cad-GFP in the membrane (lateral diffusion), which could contribute to FRAP. We then addressed which Pcdh18a domains are important for E-cad endocytosis. Full-length Pcdh18a is localized most prominently in vesicles (Figure 4B). Pcdh18a lacking the intracellular domain (Pcdh18a-ECD) is enriched at the membrane independent of the presence of E-cad. Co-expression of WT-Pcdh18a together with Pcdh18a-ECD to leads to cointernalization. Co-expression of E-cad with Pcdh18a leads to enhanced formation of intracellular E-cad positive puncta, possibly, E-cad+ endocytic vesicles, whereas Pcdh18a-ECD is unable to route E-cad to endocytic vesicles. These data suggest that the extracellular domain of Pcdh18a is important for homophilic interactions, whereas the intracellular domain of Pcdh18a is important for internalization of Pcdh18a homodimers and E-cad/Pcdh18a heterodimers. In support of this hypothesis, co-expression of E-cad and Pcdh18a-ECD reduces recovery of E-cad-GFP after bleaching (Figure 4A), suggesting that Pcdh18a-ECD reduces E-cad endocytosis and recycling. Protocadherins and cadherins also form dimers between neighbouring cells to regulate cell adhesion in a cluster [19]. Similarly, we found that Pcdh18a also forms trans-interactions with E-cad, followed by endocytosis of the E- cad/Pcdh18a plaques (Supplementary figure 6A). In summary, we conclude that Pcdh18a shifts the balance from degradation towards recycling by routing E-cad to the Rab-11 positive recycling endosomes, which leads to the stabilization of E-cad. Thus, our data suggest that Pcdh18a increases the E-cad levels in the membrane and in vesicles in the NTC cluster.

**Figure 4:**
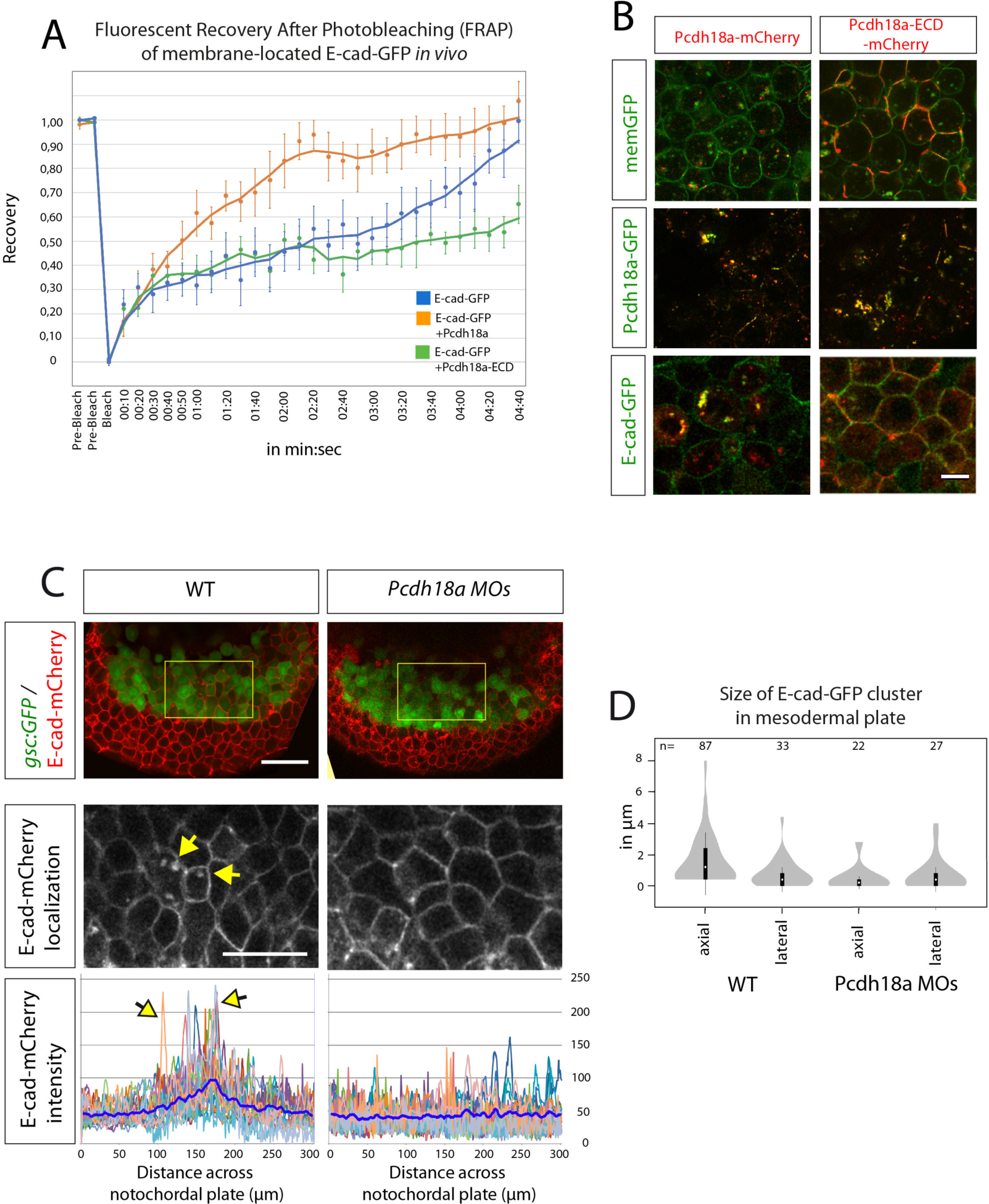
Pcdh18a domains and their importance with regard to endocytosis. A, After photobleaching of a 3µm spot at the cell membrane of E-cad-expressing embryos, new E-cad-GFP molecules moved into the bleached area from adjacent membrane regions, resulting in a return of 90% of fluorescence within 4:20min (blue). Co-expression of Pcdh18a increased the speed of recovery and a 90% recovery was reached after 2:20min (orange). Co-expression of Pcdh18a-ECD diminished FRAP of E-cad-GFP (green). Moving-average trendline was calculated with period 3. B, Confocal images of zebrafish embryos at 5hpf (50%epiboly). Embryos were injected with 0.1ng mRNA of indicated constructs. In the deletion construct Pcdh18a-ECD the intracellular domain was replaced by a mCherry. Pcdh18a- mCherry was localized to vesicles, whereas Pcdh18a-ECD-mCherrry was strongly localized to the cell membranes. Pcdh18a-GFP/Pcdh18a-mCherry and Pcdh18a-GFP/Pcdh18a-ECD-mCherry showed co-localization at the membrane and in vesicles suggesting homophilic interaction. Pcdh18a-mCherry/E-cad-GFP suggest heterophilic interaction. Pcdh18a-ECD-mCherry and E-cad-GFP were observed mainly at the membrane and did not co-localize suggesting that the intracellular domain of Pcdh18a is required for interaction and co-internalization with E-cad. Scale bar, 10 µm. C, tg(gsc:GFP) embryos were injected with 0.1 ng of the e-cad-mCherry mRNA or co-injected with the pcdh18a Morpholino (0.5 mM) and subjected to confocal microscopy analysis at 8 hpf. A cross-section at the level of the NTC (Supplementary figure 4A, section plane 2) reveals enhanced E-cad localization at the plasma membrane and in endocytic vesicles, as shown by a projection of five fluorescence intensity histograms of five different embryos. Scale bar, 100 µm. D, Bean plots shows the distribution, means, and standard deviations of the sizes of e-cad-GFP clusters in the lateral and axial mesodermal plate measured in 20 WT and Pcdh18a morphant embryos.

Next, we analysed the function of endogenous Pcdh18a with respect to E-cad trafficking in the NTC. Subcellular E-cad concentrations were measured across the central NTC of *tg(gsc:GFP)* transgenic zebrafish embryos (Supplementary figure 4A, arrow 3). We discovered enhanced localization of E-cad-mCherry in the membrane and in vesicles in the NTC, as indicated by the peaks in the E-cad intensity histograms (Figure 4C) and quantification of the number and sizes of E-cad clusters in the mesodermal plate (Figure 4D). We did not observe these focal E-cad accumulations in the LPM cells surrounding the NTC or in the NTC of Pcdh18a-deficient embryos, suggesting that NTC display enhanced endocytosis of E-cad, presumably mediated by endogenous Pcdh18a.

### Recycling of E-cad plaques determine cell migration

An increase in cadherin-mediated cell adhesion complexes usually leads to cluster formation through cell separation and reduced cell migration [20,21]. A classic example is the epithelial-mesenchymal transition, in which downregulation of E-cadherin is linked with epithelial junction disassembly and subsequent enhancement of developmental migration and malignant invasion of single cells [22]. However, our experiments suggest that enhanced Pcdh18a/E-cad presentation might increase migration of cell clusters. To resolve this discrepancy, we employed an IBIDI-based wound-healing assay to analyse the function of the NTC key players in cell migration in a controlled in vitro experiment. We ectopically expressed Pcdh18a, E-cad, and Fzd7a in highly motile human cervical cancer cells (HeLa), which display low endogenous expression of these transmembrane proteins [23]. We found no alteration in the migratory activity after transfection of Pcdh18a (Figure 5A,B). Conversely, we observed that the motility of the E-cad-transfected HeLa cells was significantly decreased. Notably, we found that the expression of Pcdh18a in the E-cad-positive cells sped up cell migration to levels comparable to the control cells. This result is in concordance with recently published data showing that the cytoplasmic sites of several δ-Pcdhs interact with WAVE complex member [24]. One example is the related zebrafish gene Pcdh18b, which controls actin assembly via Nap1 [25]. Furthermore, Pcdh8 suppresses C-cadherin-mediated cell adhesion and promotes the Wnt-PCP pathway-dependent cell motility [26]. In Xenopus, Fzd7 diminishes cis-dimerization of Cadherin (C-cadherin) and protocadherin (PAPC) and thereby weaken cadherin-mediated adhesion [27]. We found that, Fzd7a-positive HeLa cells and Pcdh18a/Fzd7a-positive HeLa cells moved into the cleft significantly faster than the control cells (Figure 5A,B). Like in Xenopus, we observed a similar reduction of migration after coexpression of E-cad and Fzd7a. We performed a triple transfection of E-cad, Pcdh18a and Fzd7a in HeLa cells to mimic the expression profile of the NTC. These cells were observed to move with significantly increased speed into the cleft as clusters of 15-20 cells (Figure 5A,B). In summary, we find that E-cad expression reduces migration of HeLa cells and Pcdh18a antagonizes the effect of E-cad. Fzd7a promotes motility, which is antagonized by E-cad. Finally, co-expression of Pcdh18a partially restores increased motility in E-cad/Fzd7a positive Hela cells, possibly because Pcdh18a antagonizes increased cell adhesion from E-cad. Together, with the data on co-localisation (Figure 3) and endocytosis (Figure 4), these observations suggest a functional interaction of Pcdh18a/E-cad/Fzd7a influencing migratory behaviour.

**Figure 5:**
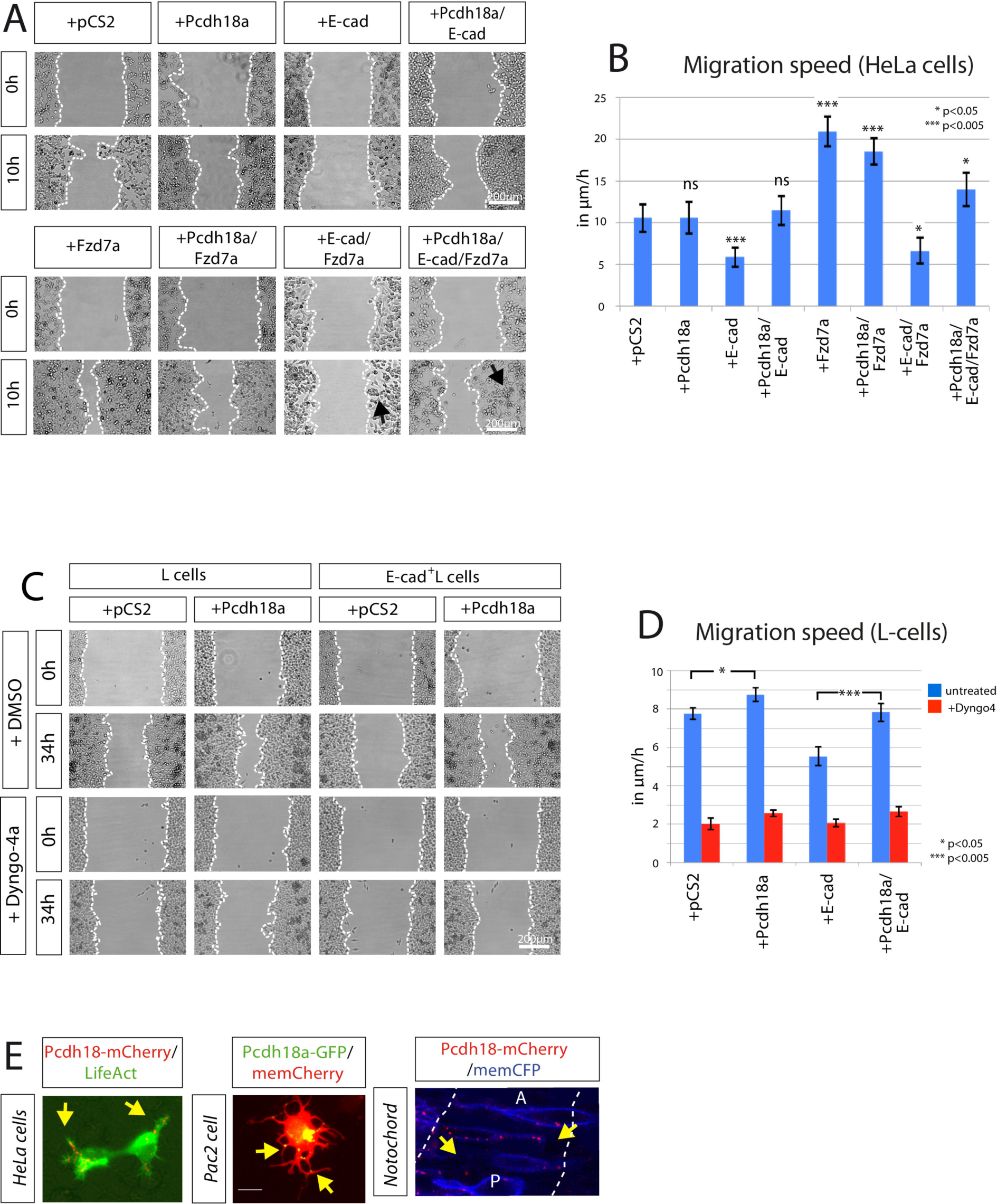
Pcdh18a affects E-cad dependent cell migration. A, Wound-healing assay in HeLa cells. Cells were triple-transfected with the indicated constructs and their migratory behaviour was monitored for 10 h after removing the insert. Arrows mark cell clusters, and the dotted line shows limits of the confluent cell layer. B, Quantification of the migration speed of HeLa cells. C, Wound-healing assay in L-cells. Cells were transfected with indicated constructs. After 24 h, L-cells were treated with DMSO (1%) or Dyngo4a endocytosis inhibitor (1 µM in DMSO 1%). The migratory behaviour of cells was monitored in a time lapse for 34 h. D, Quantification of the migration speed of L-cells after blocking endocytosis. The wound healing assays were conducted in independent triplicate, distances of the gap were measured at 10 fixed positions. Mean values, SEM and significance are indicated E, Co-transfection of Pcdh18a and LifeAct in HeLa cells revealed asymmetric, subcellular distribution (72% leading edge, 14% trailing edge, n=12). In Pac2 fibroblasts, Pcdh18a-GFP is localized in the retracting protrusions. The mosaic expression of Pcdh18a- mCherry and memCFP in the NTC shows that Pcdh18a is preferentially localized in the posterior region (a=67%, p=24%, n=8 cells in 5 embryos) of the developing zebrafish embryo at 6 hpf. Scale bar, 15 µm.

To measure cell motility with altered endocytic routing of E-cad, we set up a robust experimental procedure utilising L-cells in an IBIDI-based wound-healing assay. The transfection efficiency of Pcdh18a to L-cells was about 40% (Supplementary figure 7A). L-cells were then treated with Dyngo-4a, a blocker of dynamin-dependent endocytosis. Similar to our results with HeLa cells, we found that Pcdh18a expression increased cell migration in E-cad-GFP+ L-cells (Figure 5C,D). The reduction in endocytic trafficking of E-cad through the inhibition of Dynamin function led to a significant decrease in cell migration, which could only partially be compensated by Pcdh18a expression. To provide further evidence that Pcdh18a acts functionally together with E-cad, we compared the migration speed of E-cad-GFP+ L-cells transfected with Pcdh18a with E-cad-GFP+ L-cells treated with the E-cad blocking antibody DECMA [28]. We found that both treatments increased migration speed of E-cad-GFP+ L-cells in a similar way (Supplementary figure 7B). The in vitro experiments suggest that E-cad leads to decreased cell migration; however, Pcdh18a converts these slow migrating cells into fast migrating clusters by enhancing E-cad recycling.

E-cad adhesion plaques play a pivotal role in maintaining physical interactions and in serving as signalling platforms that control cell polarity and directed migration [29]. Furthermore, N-cad-based adhesion complexes are preferentially endocytosed in the rear half of the cells and undergo continuous intracellular retrograde transport to orchestrate collective migration [30]. To examine the subcellular localization of Pcdh18a in migrating cells we expressed Pcdh18a-mCherry and LifeAct in HeLa cells, in Pac2 zebrafish fibroblasts, and in notochord clones in the zebrafish embryos (Figure 5E). Pcdh18a-mCherry was found to be mainlineocalized to the trailing edge of HeLa cells, and we observed retrograde transport of Pcdh18a in protrusions in Pac2 cells. We further found enhanced Pcdh18a localization at the posterior membranes of notochord cells.

### Pcdh18a/E-cad positive tip cells could organize notochord organization

To study the self-organization of the mesodermal tissue during gastrulation a lattice-based Cellular Potts Model (CPM) was implemented based on the Glazier-Graner-Hogeweg Model [31]. This model recaptures our experimentally accessible cell properties and is defined by intercellular adhesion energies, intracellular surface and volume constraints, and migration properties (Figure 6A). The CPM was used to address the question of which properties of individual cell groups were required to generate intercalation of the migrating cell layer into a notochord-like structure. In our modelling framework, every cell is represented in an object-oriented fashion by a physical location in the tissue, as well as cell type-dependent physical properties such as cell-cell adhesiveness, relative migration speed, and migration direction. We specified four different cell types: the leading PPM cluster, the central NTC, the trailing notochord plate, and the surrounding LPM. Every cell, together with its interactions with its next neighbours, gives an energetic contribution to the total energy of the system, as specified in the Hamiltonian function of the CPM (see Materials and Methods). The system evolves from one state to the next by employing the Metropolis Monte-Carlo criterion for comparing these energetic contributions, giving rise to a statistically descriptive end state. The simulation started with cells ingressing from the epiblast and forming the mesodermal cell layer. The first cells to appear were the E-cad-positive PPM cells (Figure 6A). These cells are an anteriorly migrating group of cells that are characterized by high cell-cell adhesiveness. Therefore, it is unlikely that these cells change neighbours. These cells were followed by a group of anteriorly migrating NTC surrounded by LPM cells. New LPM cells pushed into the field as the cells migrate forward. Simulations were stopped when the LPM reached an extension of 400 [jm, which equates to 8hpf. We studied three different scenarios for the NTC: mobile leaders, adhesive leaders, and mobile & adhesive leaders. Based on our observations of alter cell migration in vitro (Figure 5A), we increased the cell motility μ of the mobile leading NTC by 100% compared to their neighbours (Figure 6A, mobile leaders). We observed that the NTC was seen to exhibit an oblong shape that was perpendicular to the anteroposterior axis, similar to a droplet hitting a wall. This result suggests that the differential adhesion between the PPM and the NTC, along with the weak cell-substrate interaction, inhibits the elongation of the notochordal plate. The adhesion of the NTC was then increased without altering the cell migration speed (adhesive leaders). We observed that the trailing notochordal plate did not compact and showed a remarkably similar phenotype of embryos with reduced Pcdh18a levels (Figure 1C). Interestingly, it seems that the NTC slows down the migration anterior located, medial PPM cells, leading to a concave tissue shape. Finally, we increased both the migration speed and adhesion (Figure 6A, mobile & adhesive leaders). We found that the shape of the tissue changed: the E-cad-positive leading PPM formed a convex outline and thus resembled the curved shape of the PPM in vivo. Even more striking than changes to the tissue shape, we observed that the central NTC and the notochordal plate condensed to a rod-shaped structure. We therefore hypothesize that the organization of the mesodermal plate requires the stretching force of the NTC on the notochordal plate (see phenotypical analysis in Figure 1D). As the CPM is nearest neighbour-based, we conclude that the stretching forces generated from the NTC primarily operate on the trailing notochordal plate and secondarily on the adjacent LPM cells, pulling them towards the midline.

**Figure 6:**
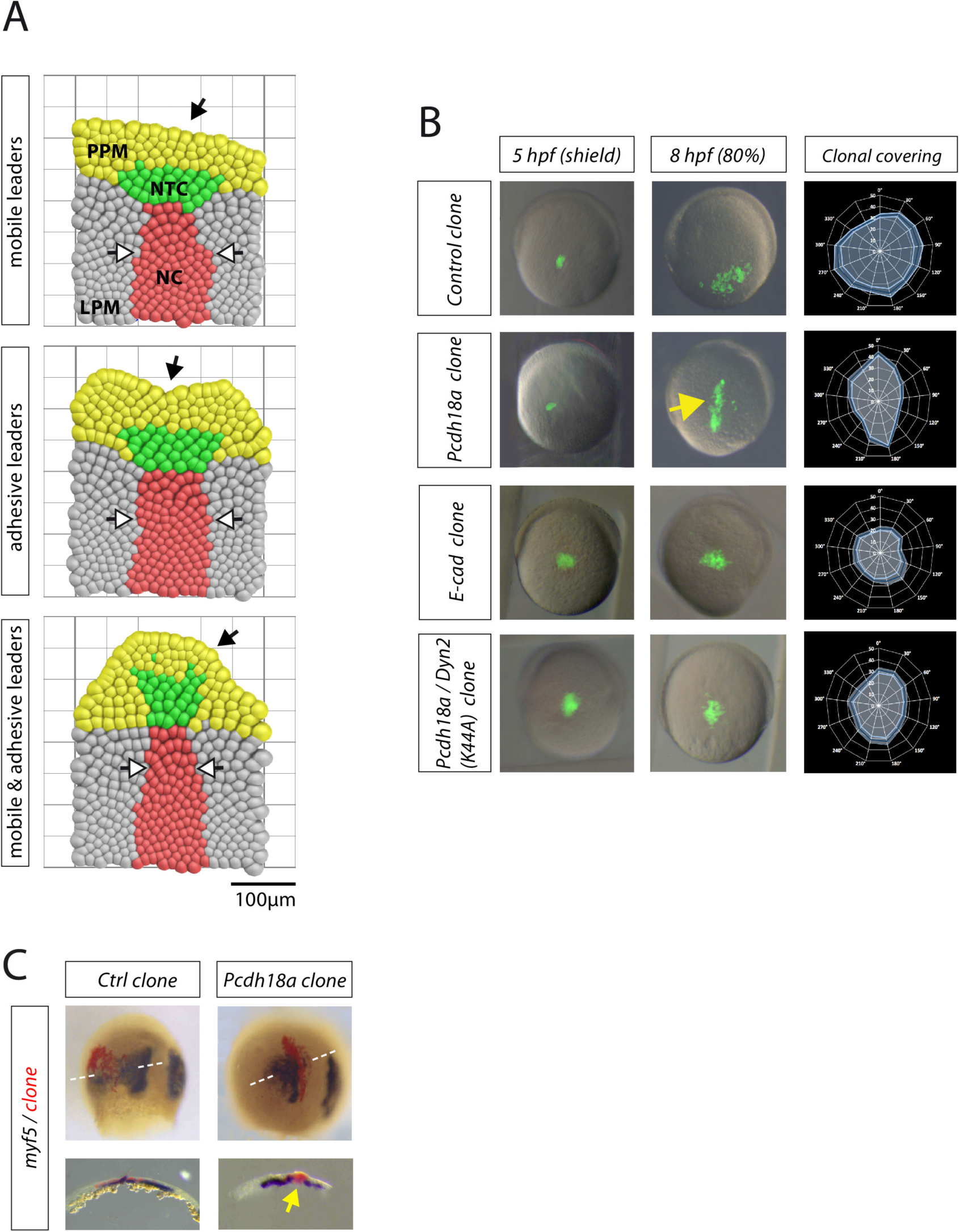
Directed cohort migration results in the formation of the rod-shaped notochord. A, Visual rendering of the results of the simulations. See Supplementary figure 8 for the description of the CPM. B, Embryos were microinjected with mRNAs for the indicated constructs (pcdh18a: 0.3 ng, e-cad : 0.4 ng, dyn2K44a : 0.2 ng). At 5 hpf, approximately 50 cells were grafted into the lateral embryonic margin of uninjected host embryos. At 8 hpf, the migration and the directionality of the cell clusters were analysed. Animal pole was set to 0°, vegetal pole was set to 180°. Blue line indicates mean value of clonal coverage measured in ten different embryos per experiment and white lines indicate SEM. C, Embryos from (B) were fixed and subjected to ISH for the LPM marker myf5. Horizontal cross-sections revealed the formation of an ectopic rod-shaped structure of the Pcdh18a-positive clones in the LPM.

Thus far, our experiments and simulations suggest that Pcdh18a expression in the NTC is required to organize the shape of a notochord either by regulating gene expression in the notochord or by controlling cellular mechanics. In a test of the first hypothesis, we found that Pcdh18a-positive clones did not induce ectopic expression of the notochord marker *ntl* in the LPM (Supplementary figure 9A), in contrast to Chordin/Dkk1 positive clones, which generate a secondary Spemann organizer followed by the induction of a secondary notochord. Next, we performed a deep mRNA sequencing analysis of Pcdh18a-overexpressing embryos and Pcdh18a-morphant embryos. In concordance with our results, we did not detect alteration in the expression of mesodermal genes such as *chordin, gsc, ntl, shh a*, and *shh b*, or in the expression of *e-cad* and *fzd7a* (Supplementary figure 9B). During collective cell migration, the Rac-1 mediated polarization of leader cells, the interaction of leader and follower cells and the migration process with retrograde flow of the adherens junctions, has been studied during oogenesis in *Drosophila*, lateral line formation in zebrafish, neural crest formation in Xenopus and cancer invasion [29]. However, the molecular mechanism regulating increased adhesiveness within a cluster and fast migration of the very same cluster on a substrate - i.e. neighbouring cells - at the same time is not well understood. Therefore, we tested the second hypothesis; whether the Pcdh18a-positive clones determine tissue shape by regulating cohesive cell migration. Cells were grafted from the lateral blastoderm margin from a Pcdh18a- positive donor into the lateral blastoderm margin of a WT host at 5hpf. The cells formed a compact and elongated cell cluster, which migrated towards the animal pole (Figure 6B). The re-organization into rod-shaped NTC was even more obvious in horizontal sections of embryos subjected to ISH for the LPM marker *myf5* (Figure 6C). We then asked whether this phenotype could be explained by enhanced E-cad localization at the cell membranes of the cluster cells, as suggested previously (Figure 3A,B). E-cad-expressing cell clones were indeed observed in clusters, however, the shape of these clusters was circular, and the clustered cells migrated slowly towards the animal pole, similar to the in vitro wound healing approach (Figure 5A,C) and the simulation ‘adhesive leaders’ (Figure 6A). Our interpretation is in accordance with recently published data suggesting that the protocadherin, PAPC, regulates endocytosis of N- cad in the anterior compartment of the forming chick somite to allow tissue re-arrangements in the forming somites [32]. Furthermore, analysis of E-cad localization during zebrafish epiboly suggests that endocytic E-cad treadmilling allowed the blastoderm cells to dynamically remodel their cell-cell contacts, leading to enhanced mobility [33]. To test our hypothesis whether Pcdh18a-regulated E-cad endocytosis promoted cohesion and migration, we grafted cells into a WT host embryo that expressed Pcdh18a and a mutated form of Dynamin2^K44A^ to block endocytosis (Figure 6B). We found that Pcdh18a/Dyn2^K44A^ positive cells form a cluster. However, similar to the E-cad-positive cell cluster, they rarely migrated towards the animal pole. Based on these results, we conclude that Pcdh18a regulates directed cohort migration behavior in cell clusters. This cell cluster may influence the intercalation of the trailing cell sheet into a rod-shaped tissue. We suggest that this process serves as an equivalent to the NTC during notochord formation in the mesodermal cell sheet.

## Discussion

### Notogenesis in zebrafish

Here we identified Pcdh18a as a regulator of tissue morphology. Pcdh18a is required for formation of the notochord in the zebrafish mesodermal plate without altering the transcriptional signature of the cells (supplementary figure 9B). Based on our data, we define three consecutive steps in zebrafish notogenesis (Figure 7). The first involuting cells are the E-cad-positive marginal cells of the PPM. Collective and directional movement towards the anterior pole is intrinsically regulated by E-cad and Wnt-PCP signalling [34]. The PPM organizes in an elongated cell cluster and moves anteriorly due to a cadherin-mediated pulling force from the posterior side [35]. Second, we identified the NTC as an axial mesodermal cell population, which forms an elongated cell cluster after ingression (Figure 7, left). In mice, a similar condensation of dispersed mesodermal cells located anterior to the forming Spemann organizer was observed in the early gastrula and these cells give rise to the anterior part of the notochord [36]. We observed that the NTC marched towards the animal pole as a cohesive cell group during gastrulation (Figure 7, right). thus, the NTC pushed the anteriorly located PPM ahead, leading to the formation of a convexly curved mesodermal cell sheet. There is indeed supporting evidence that a leading-edge mesodermal cell group in *Xenopus* migrates faster than the axial mesodermal cells to exert a pushing force on the PPM [37]. Third, the NTC guide the notochord plate cells, to intercalate and elongate either by pulling forces or by signalling. We conclude that this process leads to the condensation of the trailing notochord.

**Figure 7:**
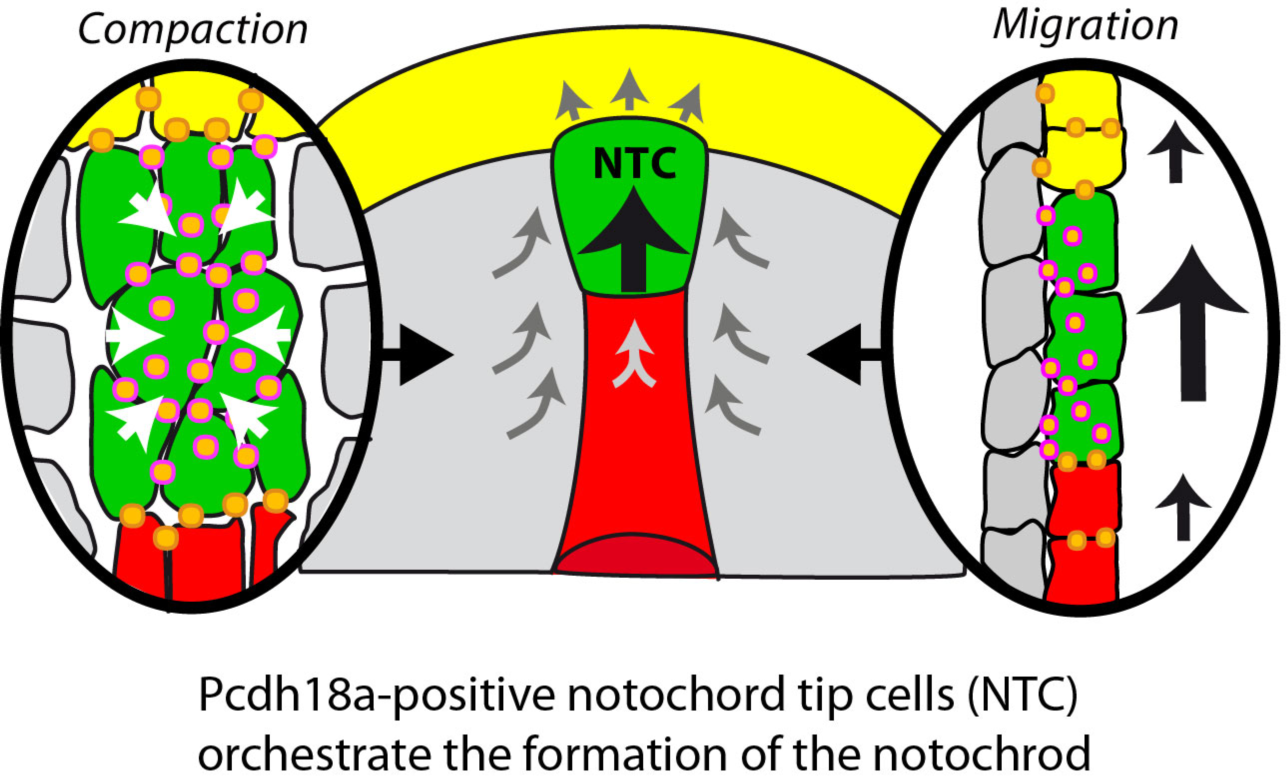
Schematic summary of the function of the NTC in notochord formation. Pcdh18a/E-cadherin adhesion complexes (orange dots) increase cell adhesion within the NTC, leading to the cluster formation (left). In parallel, Pcdh18a controls endocytosis of E-cadherin adhesion complexes to allow fast cohort migration of the NTC cluster (right), leading presumably to simultaneous pushing forces on PPM and pulling forces on the notochord.

### Tip cells organise notochord elongation and condensation

How does this interpretation fit with the common hypothesis for notochord formation? One hypothesis suggests that the notochordal plate cells actively migrate towards the animal pole and thereby extend their shape in the anteroposterior direction [38,39]. Indeed, time-lapse analysis in zebrafish revealed that the notochordal plate lengthen rapidly during gastrulation [40]. Interestingly, we find that the blockage of Pcdh18a function (Figure 1D,E Figure 2B) and the simulations (Figure 6A) indicate that the inhibition of NTC migration results in a halt in the condensation of the notochordal plate and inhibition of its extension. Furthermore, clonal expression of Pcdh18a lead to the formation of an ectopic rod-shape structure in the LPM (Figure 6B,C). Therefore, we argue that although the notochord cells may have the intrinsic capacity to intercalate and elongate, the NTC are required to push against the resistance of the PPM [34]. Only then, the compressive force generated by the notochordal plate cells is released along the anteroposterior axis, leading to the elongation, alignment, and intercalation of the notochord cells. Although our data does not exclude the possibility that the Pcdh18a positive NTC cells present a signal for the notochord cells to direct the cellular rearrangement in the trailing central mesoderm, we suggest that tension is the active agent. An additional hypothesis suggests that the two mesodermal wings push the notochordal plate and help to generate the rod-shaped structure of the notochord [41]. Our data suggest that blockade of Pcdh18a function uncouples partially the cell migration events in the axial mesoderm from the events in the LPM (Figure 1D; supplementary figure 4B,C). Furthermore, our simulations suggest that the condensation of the notochord facilitates movement of the LPM cells towards the midline. Consistent with this observation, there is no obvious convergent extension movements in the LPM in mice [42,43] and axial notochord progenitors still undergo convergent extension movements to trigger notochord elongation by cell intercalations [36,44].

### Molecular interaction of Pcdhs to regulate cell migration

The δ2 subfamily comprises five members - Pcdh8, Pcdh10, Pcdh17, Pcdh18 and Pcdh19 - reviewed in [16]. The function of Pcdh8 (also known as PAPC) in early development has been intensively studied in *Xenopus* and zebrafish. In *Xenopus* embryos, Pcdh8 is expressed in the paraxial mesoderm and is involved in cell sorting and convergent extension during gastrulation [45]. Pcdh8 seems not to act as an adhesion molecule, however, its ectopic expression promotes cell sorting in the lateral plate mesoderm [46,47]. Molecular analysis suggests that Pcdh8 acts in concert with C-cadherin. Pcdh8 forms complexes with Fzd7/Wnt11 to suppress C-cadherin-mediated cell adhesion function [27,48]. Regulation of early morphogenesis through the δ2- group protocadherins has also been reported in zebrafish. Knockdown of *Pcdh18a* by using morpholino oligonucleotides causes impaired cell movements [12], and *Pcdh19* knockdown results in impaired convergence during neurulation [49]. Our observations suggest that Pcdh18a interacts with E-cad and Fzd7a endocytosis and thus remodels cell migration and adhesion. A possible molecular mechanism has been recently suggested that Pcdhs - including Pcdh18a - accelerates cohort tissue movement by actin cytoskeleton reorganization through the WAVE complex [24,50]. Cadherin adhesion plaques respond to mechanical pulling forces by eliciting a strain-stiffening response [51]. Therefore, it is tempting to speculate that the induction of tension on the forming notochord by the NTC leads to a reinforcement of an axial structural scaffold. Our data suggest that Pcdh18a-mediated E-cad turnover in the notochord tip cells generates forces that are required for the organization of the mesoderm and generation of the hallmark of Chordata - the notochord.

## Materials and Methods

### Zebrafish husbandry

Adult zebrafish (Danio rerio) were maintained at 28.5 °C on a 14 h light/10 h dark cycle [52]. The data we present in this study were acquired from an analysis of wild-type zebrafish (AB) and transgenic zebrafish lines: tg(-1.8 gsc:GFP)ml1 [34], tg(-2.2shha-:GFP:ABC)sb15Tg [53] tg(h2afx:EGFP:rab5c)mw5Tg, tg(h2afx:EGFPrab7)mw7Tg and tg(h2afx:EGFPrab11a)mw6Tg [54].

### Functional analysis

Transient knockdown of gene expression was performed using Morpholino oligonucleotides (MO). The following antisense oligomers were used:

Control MO 5’-CGAAGTCTACGTCGGAATGCAGG-3’ [55];

Pcdh18a UTR MO 5’-TCCGTCAGGCACTGCAAAAATATAC-3’ [12];

Pcdh18a Translation Blocking MO 5’-ACCCTTGCTAGTCTCCATGTTGGGC-3’ [12]. Pcdh18a function was inhibited by injecting a 0.5 mM concentration of each Morpholino oligomer at the one-cell stage. In parallel, we used the CRISPR/Cas9 system to target the Pcdh18a locus. All sgRNAs were designed and evaluated for potential off-target sites using CCTop, the CRISPR/Cas9 target online predictor (http://crispr.cos.uni-heidelberg.de/) [56], selecting the target sequence 5’ - CAGAGCAAGTTTGAGTAAAGTGG - 3’. For sgRNA assembly, a pair of synthesized oligomers (5’ - TAGGGAGCAAGTTTGAGTAAAG - 3’; 5’ - AAACCTTTACTCAAACTTGCTC - 3’) was annealed, ligated into the DR274 (Addgene plasmid #42250) vector[57], linearized with FastDigest Eco31l (Thermo Fisher Scientific), and used for in vitro transcription with the T7 MegaShortScript Kit (Ambion). The Cas9 protein was purchased from Thermo Fisher Scientific (B25640). Prior to the injection, the Cas9 protein and sgRNA were diluted in RNAse-free water and incubated for 5 min at RT for complex formation [13]. For the identification the following primers were used: pcdh18a for:

TGGCACTAAAGGAGGCTTTG; pcd18 rev mut/WT: CACTTTACTCAAACTTGCTCTGC; pcdh18a rev ctrl: ACCAGGATGGAGAGATCAGC.

The following plasmids were used for the overexpression studies: Pcdh18a in pCS2+, Pcdh18a-GFP in pCS2+, Pcdh18a-mCherry in pCS2+, GPI-anchored mCherry (memCherry) in pCS2+, GAP43-GFP, Frizzled7-CFP (gift from Dietmar Gradl, KIT), and E-cadherin-GFP and E-cadherin-mCherry (gift from Erez Raz, University of Münster). Capped and in vitro transcribed mRNAs (mMessage Machine Kit, Ambion) were microinjected into one-cell stage embryos.

### Preparation of the genomic DNA and PAGE analysis

After microinjection, individual embryos (uninjected control embryos and CRISPR transient mutants) were processed for genomic DNA extraction [58]. The standard PCR conditions were: 94°C for 5 min; 94°C for 30 s, 52°C for 25 s, 72°C 30 s for 35 cycles; and 72°C for 5 min, followed by denaturation for 3 min at 95°C. The PCR products were resolved by electrophoresis on non-denaturing polyacrylamide gels containing 15% acrylamide, 1X Tris-EDTA (TAE), 10% ammonium persulfate, and TEMED. After 2 h of electrophoresis at 120 V and 400 mA, the polyacrylamide gel was immersed in a 0.5% ethidium bromide solution for 10 min and then visualised.

### Compounds and inhibitors

Dechorionated embryos from the sphere stage to 80% epiboly were treated with 30 [jM SB505124 (Sigma) to block mesoderm formation. In cell culture, 1 µM Dyngo-4a (Tocris) was used to block clathrin-mediated endocytosis.

### Cell culture experiments

HeLa cells were obtained from Sigma. L-cells that were stably transfected with human E-cadherin-GFP (E-cad-GFP+ L-cells) were provided by Clemens Franz (KIT), [59]. Both cell types were cultured in high-glucose DMEM, supplemented with 10% FBS and 1% Pen/Strep. The cells were transfected using the FuGene HD Transfection Reagent (Invitrogen) at 80% confluence.

### Wound healing assay

Wound healing assays were performed using cell culture inserts (IBIDI). Approximately 5 x 104 cells were cultured in each cell culture reservoir, which was separated by a 500 µm thick wall. After 6 hours of cultivation, the culture inserts were removed and cell migration was monitored for several hours using an Axiovert 800 M inverted microscope. The obtained timelapse images were analysed using ImageJ software (NIH). To block E-cadherin mediated adhesion by adherens junctions, E-cad-GFP+ L-cells were treated with 50mg/ml of the anti-E- cad antibody DECMA-1 (U3254, Sigma Aldrich) during the wound healing assay.

### Western blotting

For the Western blots, whole-cell extracts and whole embryo extracts were prepared and resolved by 2-10% gradient SDS-PAGE. The proteins were then transferred to a PVDF membrane. The membrane was incubated with anti-GFP (Sigma, 1:1,000) and anti-PCNA (Abcam, 1:5,000) primary antibodies for 4 hours at room temperature.

### Fluorescence Recovery After Bleaching (FRAP) Assay

For the FRAP assay, 4 embryos were micro-injected with either E-cad-GFP (100ng mRNA), with E-cad-GFP (100ng mRNA)/ Pcdh18a-mCherry (200ng mRNA), or with E-cad-GFP (100ng mRNA)/ Pcdh18a ECD (200ng mRNA). After 5hpf, embryos were mounted in agarose. A laser light (405nm) bleached 3-micron spots on the cell membrane. The fluorescence within 3 independent bleached spots per embryo were tracked by confocal microscopy before and after bleaching in 10sec intervals. As E-cad-GFP moves into the bleached area, fluorescence recovers with an exponential time course. Within the spot, pre-bleaching fluorescence was set to 1 and post-bleaching fluorescence to 0.

### Three-dimensional optic flow tissue analysis

We used a custom three-dimensional optic flow code written in Matlab to measure the threedimensional velocity fields in both the ectoderm and mesoderm tissues. The memCherry expression in the ectoderm (red signal) was used to calculate ectoderm velocity and the *gsc:GFP* expression in the axial mesoderm (green signal) to calculate mesoderm velocity. To avoid overlap between the two tissues velocity fields, we manually masked the ectoderm for mesoderm measurements and the mesoderm for ectoderm measurements. A sliding box of 20*20*8 pixels was used to calculate the local velocity field in each voxel.

### Deep RNA sequencing and data analysis

Wild-type embryos were microinjected with 200 ng of Pcdh18a mRNA or 0.5 mM Pcdh18a MO. At 24 hpf, pools of 50 embryos from the wild-type and the injected clutches were collected. Total RNA extraction was performed with Trizol (Invitrogen) according to the manufacturer’s protocol. The extracted total RNA samples were tested on RNA nanochips (Bioanalyzer 2100, Agilent) for degradation. Sequencing libraries were generated with the TruSeq mRNA kit v.2 (Illumina). The size and concentration of the sequencing libraries were determined with DNA-chip (Bioanalyzer 2100, Agilent). Multiplexed samples were loaded on a total of six sequencing lanes. Paired end reads (2 × 50 nucleotides) were obtained on a Hiseq1000 using SBS v3 kits (Illumina). The sequencing resulted in 300 million pairs of 50-nucleotide-long reads. The reads were mapped against the zebrafish genome (Zv9) using TopHat version 1.4.1. Gene expression was determined with HTSeq version 0.5.3p3.

### Embryological manipulation assay

Donor embryos were microinjected with 0.5 mM control MO or Pcdh18a MO mixed with a lineage marker (miniEmerald, Thermo Fisher) at the one-cell stage. The host embryos were wild-type embryos or embryos that had been injected with Pcdh18a MO at the one-cell stage. At the shield stage, 50 cells were removed from each donor embryo with a needle and injected into the centre of the shield mesoderm of the host embryos. The host embryos were then allowed to develop until 90% epiboly (9 hpf) and fixed for ISH.

In vivo two-photon laser-targeted ablation of individual cell rows in the axial mesoderm was performed with a Leica Sp2. At 7 hpf, gsc:GFP embryos were mounted dorsal side up and ultrashort laser pulses were used to ablate the 5th GFP-positive cell row or the 15th GFP- positive cell row (indicated in supplementary Figure 3A, plane 1 & 2). The embryos were raised until 10 hpf for fluorescence imaging and subjected to ISH.

### In situ hybridization (ISH)

Prior to staining, embryos at the desired stage were fixed in 4% paraformaldehyde/PBS overnight at 4°C. Whole-mount mRNA ISH was performed as previously described [60]. Antisense RNA probes against ntl, hgg, pcdh18a, gsc, fzd7a, wnt11, dkk, chordin, snailla, snaill b and e-cadherin (cdh1) were used.

### Live embryo imaging and image analysis

Confocal image stacks were obtained using the Leica TCS SP5 X confocal laser-scanning microscope. We collected a series of optical planes (z-stacks) to reconstruct the imaged area. The step size of the acquired z-stack was 1 µm and was chosen based on the optimal z-resolution of the 63× objective with a numerical aperture of 0.9. The images were further processed using Imaris software 7.5 (Bitplane AG). Cell shape (roundness) was measured using ImageJ software (National Institutes of Health) and approximately 100 cells from each group were analysed.

### Hamiltonian for Cellular Potts Model

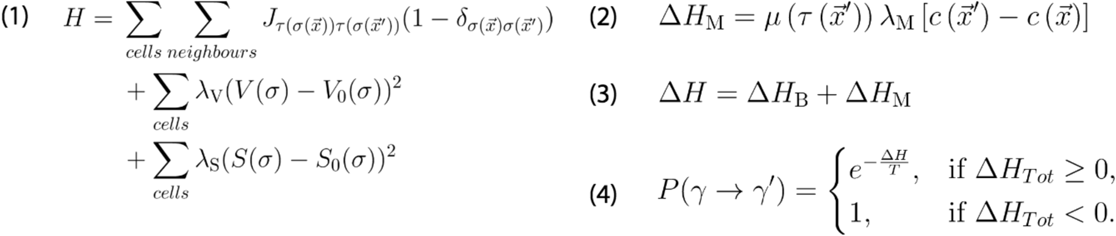

Definition for the Hamiltonian H and modifications to ΔH for our CPM. Exact parameter values are found in supplementary figure 8H. (1) The Hamiltonian for state γ consists of three sums and defines the total energy of the system. The first sum is running over each cell σ at a lattice site 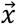 and its neighbouring lattice sites 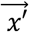. This sum represents the energetic contributions by the cellular adhesiveness J, which is dependent on the cellular types 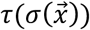and 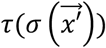.

For the exact values of J, see supplementary figure 8G. The Kronecker-delta δ makes sure that only different cells are contributing and self-interactions are excluded. The second and third sums are running over each cell, and sum up volume and surface contributions scaled by a factor λV, respectively λS. Each cell tries to preserve its original volume V0 and surface S0. (2) We defined a modification ΔHM to introduce mobility into our simulations with a linear anterior-posterior potential c. This modification is coupled to the mobility μ of the source cell at the neighbouring lattice site 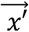 for the hypothetical state γ’ and the mobility constant λM. See supplementary figure 8A for a visual explanation of the simulation scheme. The total change in ΔH is obtained from both contributions of ΔHB = H(γ’) - H(y) and ΔHM. (3) The hypothetical state γ’ is accepted with a probability as given by the Metropolis criterion.

## Authors contributions

BB, KS, SW, and SS performed all the experiments. CS and AS generated the CPM-based simulation. VG and US performed and analysed the deep sequencing experiment. ST and TES performed and interpreted the 3D optic flow analysis. TT and JW provided support for the CRISPR analysis. BB and SS designed the experiments and wrote the manuscript.

## Acknowledgements

We would like to thank Dietmar Gradl, Keith Joung, Pierre McCrea, Angela Nieto, Erez Raz, and Máté Varga for providing the plasmids; Nicolas David and Brian Link for providing the fish lines and the Exeter BioImaging Suite for support with the FRAP experiments. We would also like to thank Máté Varga and Sepand Rastegar for providing comments on the manuscript. This project was supported by the DAAD (BB); the DFG, SCHO847-5, the LSI Exeter start-up grant (awarded to SS); and the Impuls- and Vernetzungsfond of the Helmholtz Association (awarded to AS).

## Legends supplementary figures

**supplementary figure 1.**
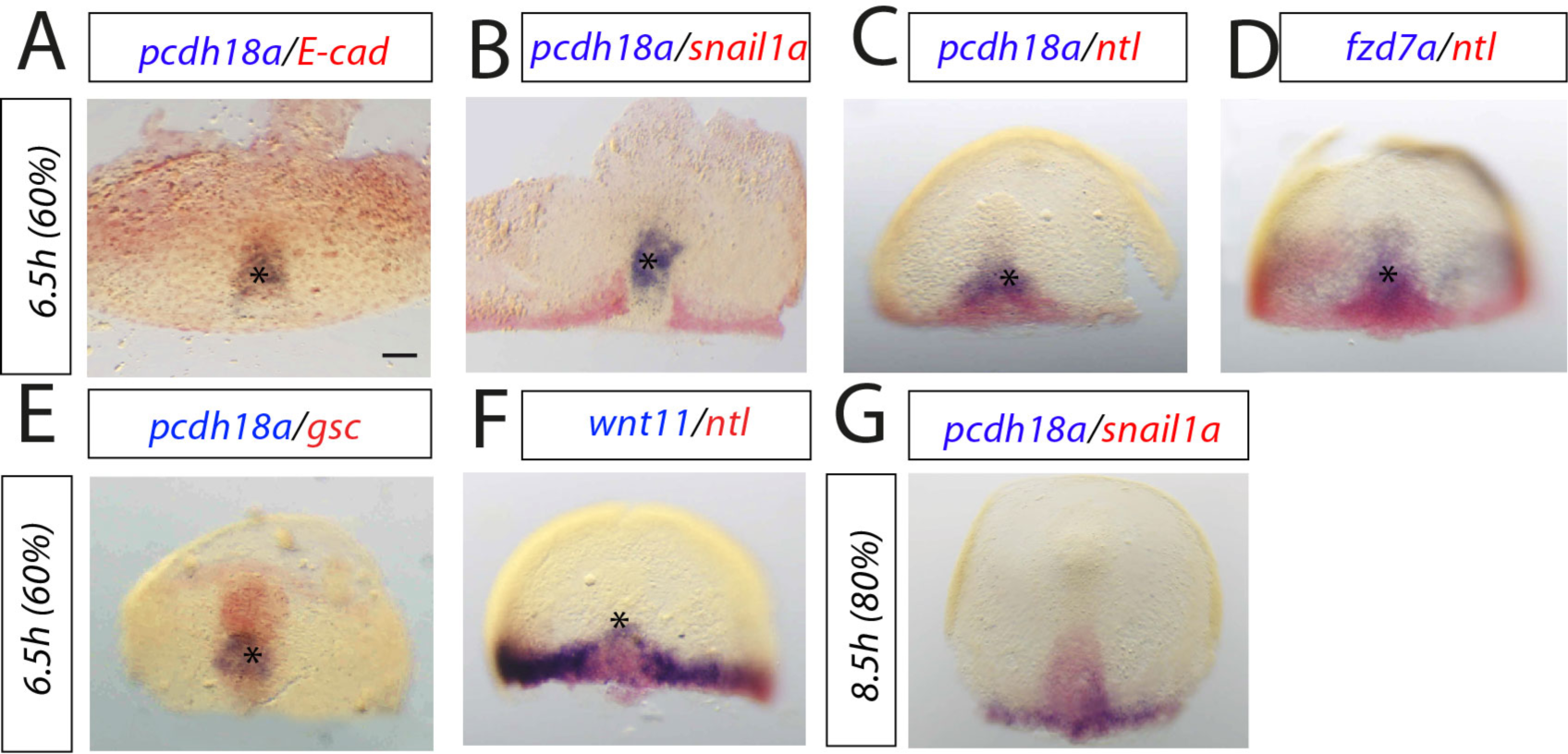
Whole-mount in situ hybridization (ISH) of zebrafish embryos. A-G, Mapping of expression patterns of indicated constructs in the mesodermal plate at 6,5hpf and 8,5hpf. Asterisks mark the notochord tip cells (NTC). Scale bar 100µm.

**supplementary figure 2.**
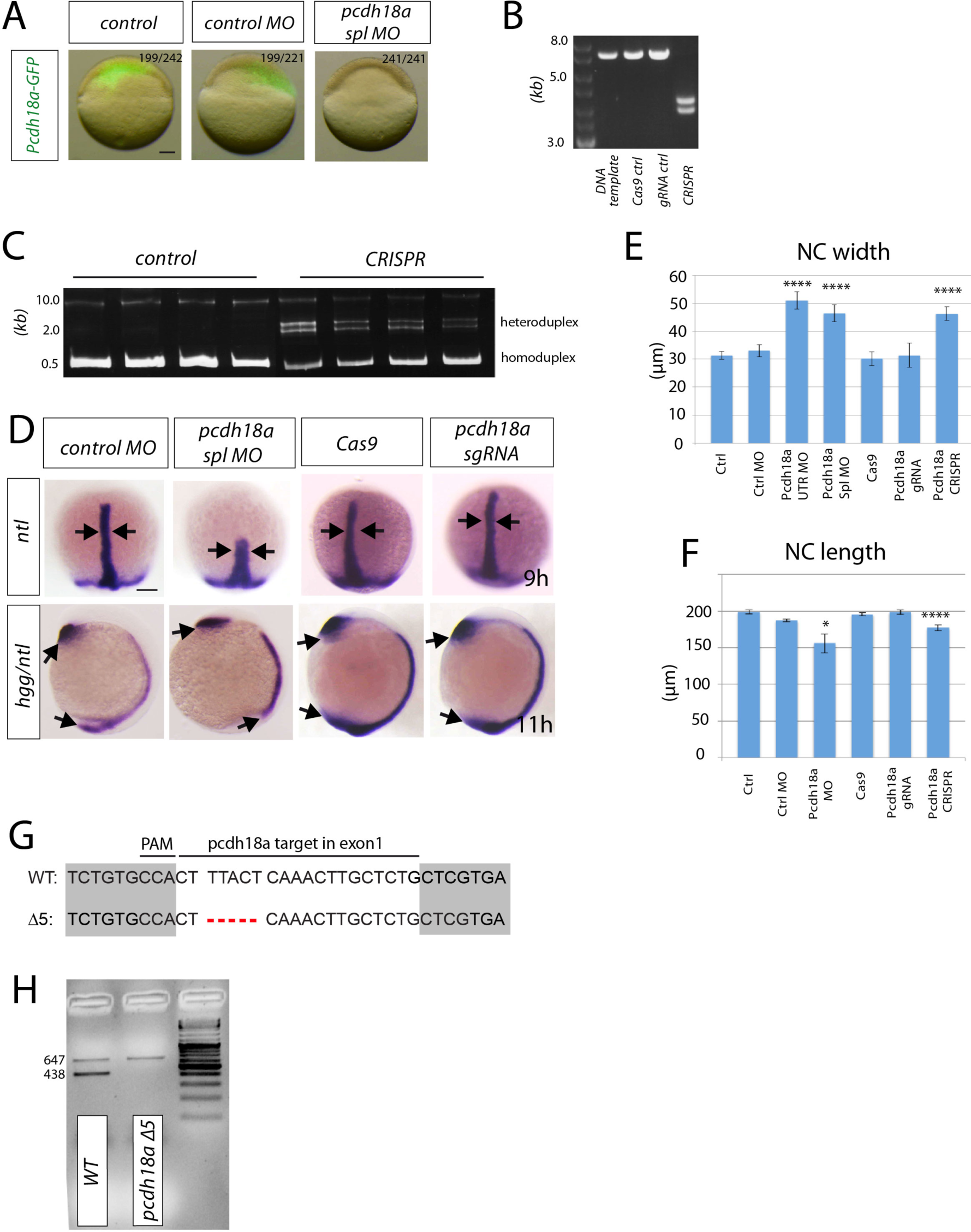
Validation of tools to inhibit Pcdh18a function in zebrafish embryos. A, Specificity of the translation-blocking Pcdh18a MO in zebrafish embryos. Embryos were microinjected in the one-cell stage with pcdh18a-GFP mRNA and control MO or MO against pcdh18a. Fluorescent images show example embryos at 5hpf. B, in vitro analysis of pcdh18a guide RNA (gRNA) efficacy. Pcdh18a-GFP containing plasmid was incubated with indicated construct. Only incubation with CRISPR (Cas9+ gRNA) leads to digest of DNA template C, PAGE-analysis provided evidence that the specific gRNA generates mutations in the endogenous pcdh18a locus in zebrafish embryos. Homo- and heteroduplexes of PCR products isolated from individual control embryos and CRISPR/Cas9 injected embryos analysed on nondenaturing acrylamide-gel. D, Embryos were injected with 0.5mM Ctrl MO, Pcdh18a MO, Cas9 protein and pcdh18a sgRNA&Cas9, raised to indicated stages and subjected for ISH for the indicated markers. Arrows mark the width of the notochord and the length of the anteroposterior axis. Scale bar 100µm. E,F, Quantifications of the notochord width and length, respectively, at 9hpf and 11 hpf, Pcdh18a MO1, Pcdh18a MO2 and Pcdh18a CRISPant embryos developed a significantly wider notochord compared to the control embryos (Ctrl: 30µm, MO1: 51µm, MO2: 46 [jm, CRISPR: 45µm) and a significantly shorter notochord compared to the control embryos (Ctrl: 200µm, MO1: 156µm, CRISPR: 177µm). *- 0.05, ****- 0.001. G, H, The 5bp deletion of the pcdh18a mutant embryos shown in Figure 1E, was confirmed by a PCR-based identification using a reverse primer spanning the deletion (product size: 438bp) and a control reverse primer located further 3’ downstream (product size: 647bp). Only *pcdh18aWT* leads to amplification of both products, whereas *pcdh18a Δ5* leads to the amplification of only one product.

**supplementary figure 3.**
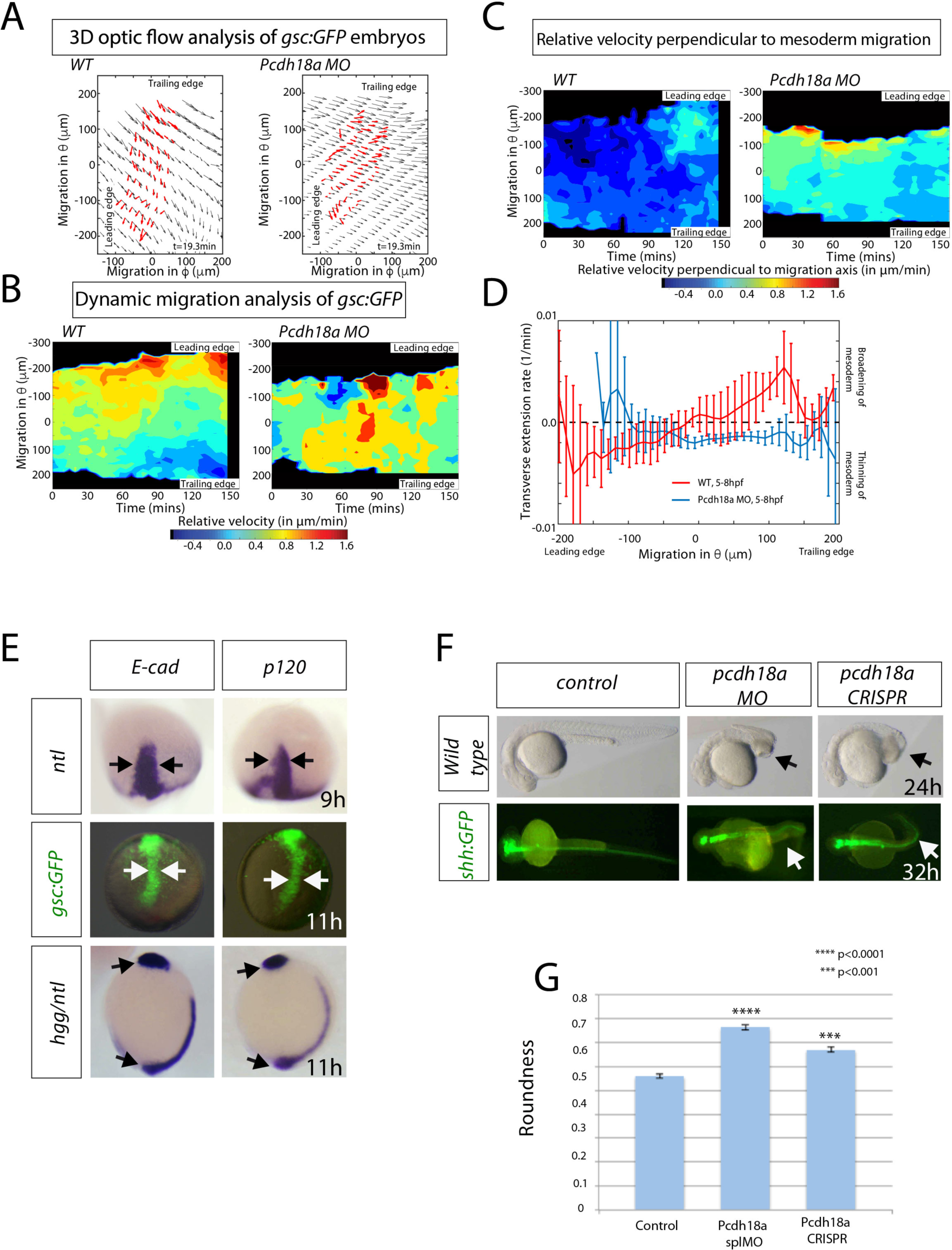
Phenotypic analysis of zebrafish embryos. A-D, 3D-Optic Flow Analysis A-B, Examples of one more WT and Pcdh18a MO embryo - colour code and nomenclature same as Figure 2B-C. C, The relative velocity (ectoderm velocity - mesoderm velocity) in the φ direction (same colour coding as Figure 2C). D, The transverse extension rate of the mesoderm for a WT and Pcdh18a MO embryo averaged over 5-8 hpf. Positive values correspond to broadening of the mesoderm perpendicular to the migration axis. Negative values correspond to thinning of the mesoderm perpendicual to the migration axis. E, ISH against ntl in 0.1ng mRNA of e-cad or p120-catenin injected embryos at 9 hpf. Arrows indicate the width of the notochord. Analysis of tg(gsc:GFP) embryos at 11 hpf, injected in the one-cell stage with 0.1ng of mRNA for e-cad or p120-catenin. Arrows indicate the width of the notochord. ISH against hgg/ntl in E-cad or p120-catenin injected embryos at 11 hpf. Arrows indicate the length body axis. F, Pcdh18a morphant embryos and CRISPant embryos display severe malformations in the trunk and tail area at 24 hpf (arrows), whereas the head seems unaltered. Analysis of tg(shh:GFP) embryos also revealed a severe bending in Pcdh18a deficient embryos (arrows). G, Quantification of cell roundness. Circularity was measured in total of 1000 cells in 5 different embryos of each tested construct. A circle has a circularity of 1.0, while noncircular shapes have a lower value of circularity. The error bars represent the SEM and significance, as indicated.

**supplementary figure 4.**
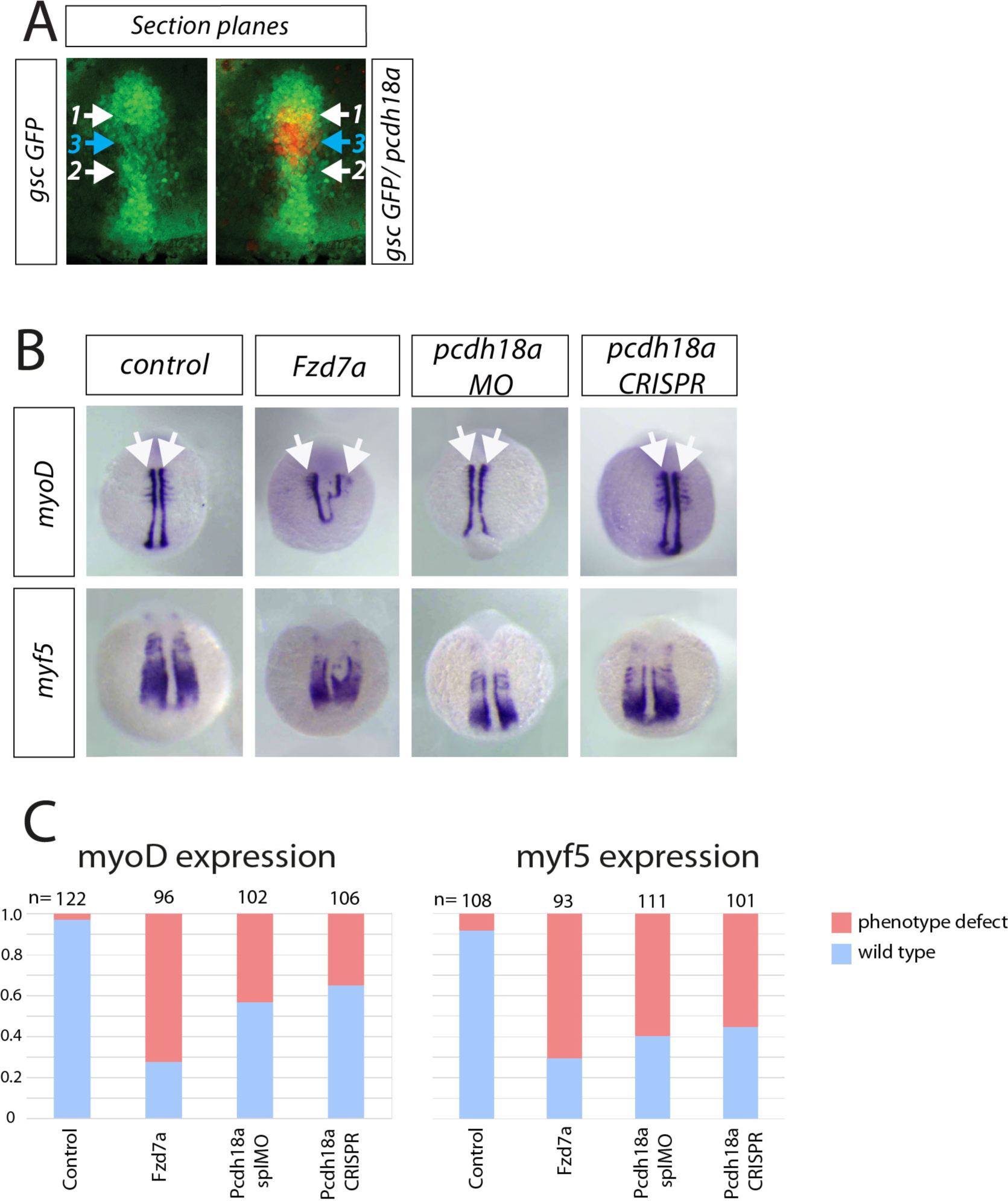
Analysis of Pcdh18a function in the LPM. A, ISH against pcdh18a followed by an antibody staining against GFP in *tg(gsc:GFP).* Arrows (1&2) indicate the sites of ablation regarding Figure 2G and arrow 3 indicate plane of image-based E-cad analysis regarding Figure 4C. B, ISH against lateral plate mesoderm marker myf5 and somite marker myoD at 12hpf. Embryos were microinjected with indicated constructs. Arrows indicate the width of the notochord. C, Phenotype classification based on *myoD* and *myf5* expressions.

**supplementary figure 5.**
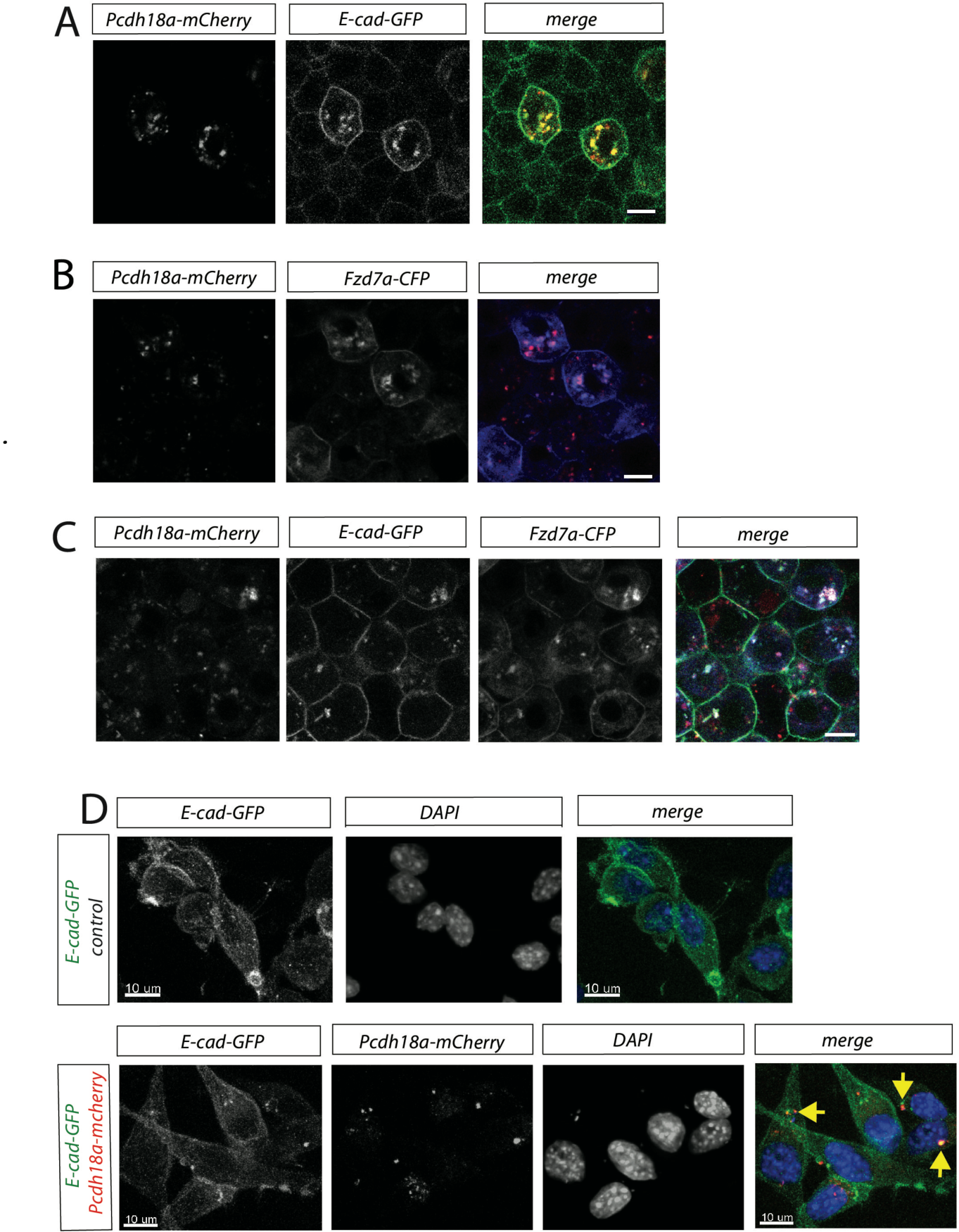
Subcellular localization of Pcdh18a and its influence on E-cad and Fzd7a. A-C, Confocal image analysis of living zebrafish embryos at 5hpf of indicated markers. D, L-cells stably expressing E-cad-GFP were transfected with Pcdh18a-mCherry and fixed, then stained with DAPI. Arrows show the co-localization of E-cad-GFP with Pcdh18a-mCherry. Scale bar 10 µm.

**supplementary figure 6.**
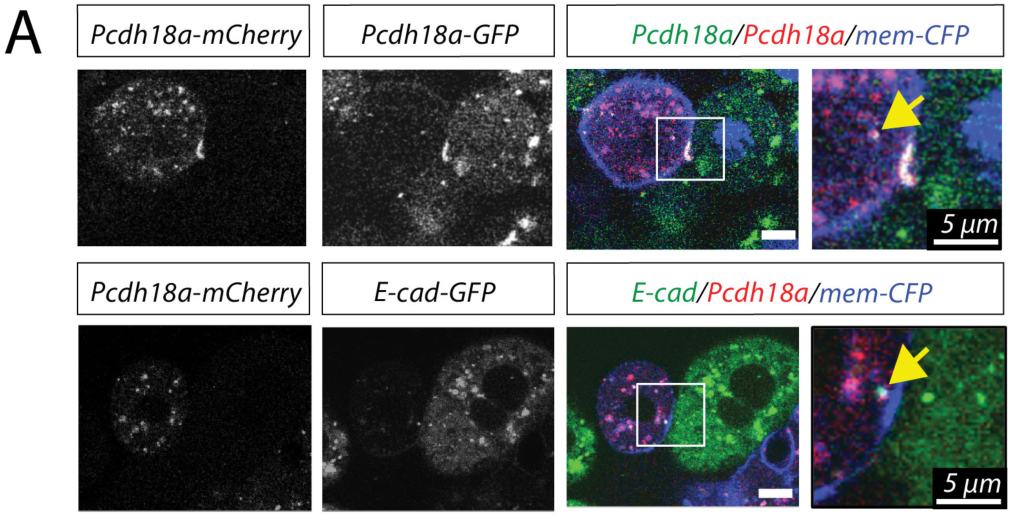
Analysis of trans-internalisation of Pcdh18a/E-cad. A, Live imaging of neighbouring cell clones expressing Pcdh18a-GFP/His-CFP or E-cad-GFP/His-CFP and Pcdh18a-mCherry/memCFP. At the 8-cell stage, the embryos were injected with 0.1 ng of pcdh18a-mCherry / memCFP mRNAs in one blastomere and e-cad-GFP or pcdh18a-GFP mRNA in the adjacent blastomere. Trans-internalization of Pcdh18a/Pcd18a and Pcdh18a/E-cad was observed (yellow arrows). Inset shows co-localization at higher magnification. Scale bars 10 µm

**supplementary figure 7.**
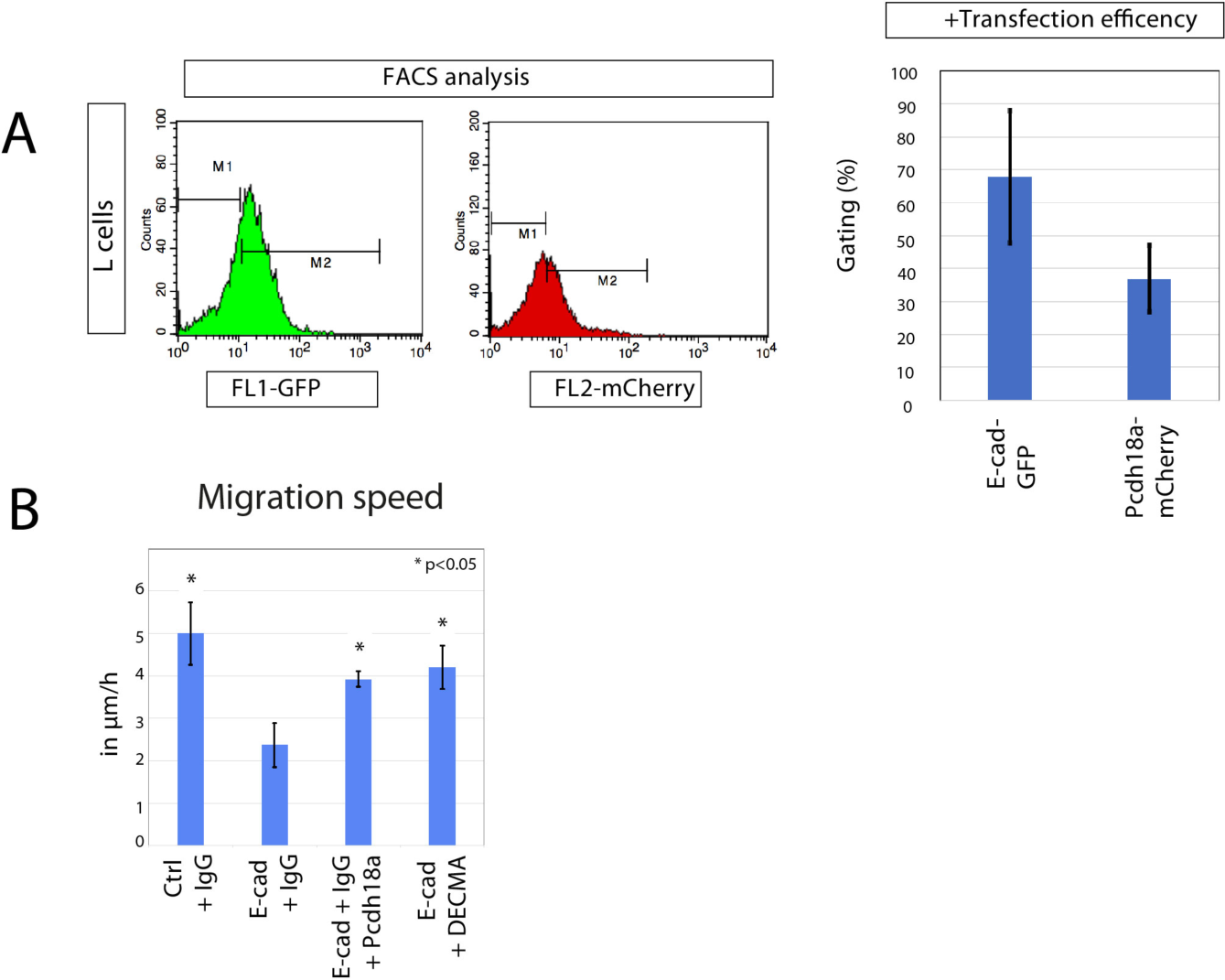
Analysis of transfection rate and migratory behaviour of L-cells. FACS analysis of L-cells transfected with E-cad-GFP and Pcdh18a-mCherry. Transfection efficiency of L-cells was measured using Fluorescence Activated Cell Sorting (FACS). M1 represents auto-fluorescence, M2 represents fluorescence of transfected constructs. B, Quantification of the migration speed of L-cells after blocking E-cad function with E-cad-blocking antibody (DECMA-1). The migratory behaviour of the cells was monitored in time lapse for 12 h. The experiments were conducted in independent duplicates. Mean values, SEM and significance are indicated.

**supplementary figure 8.**
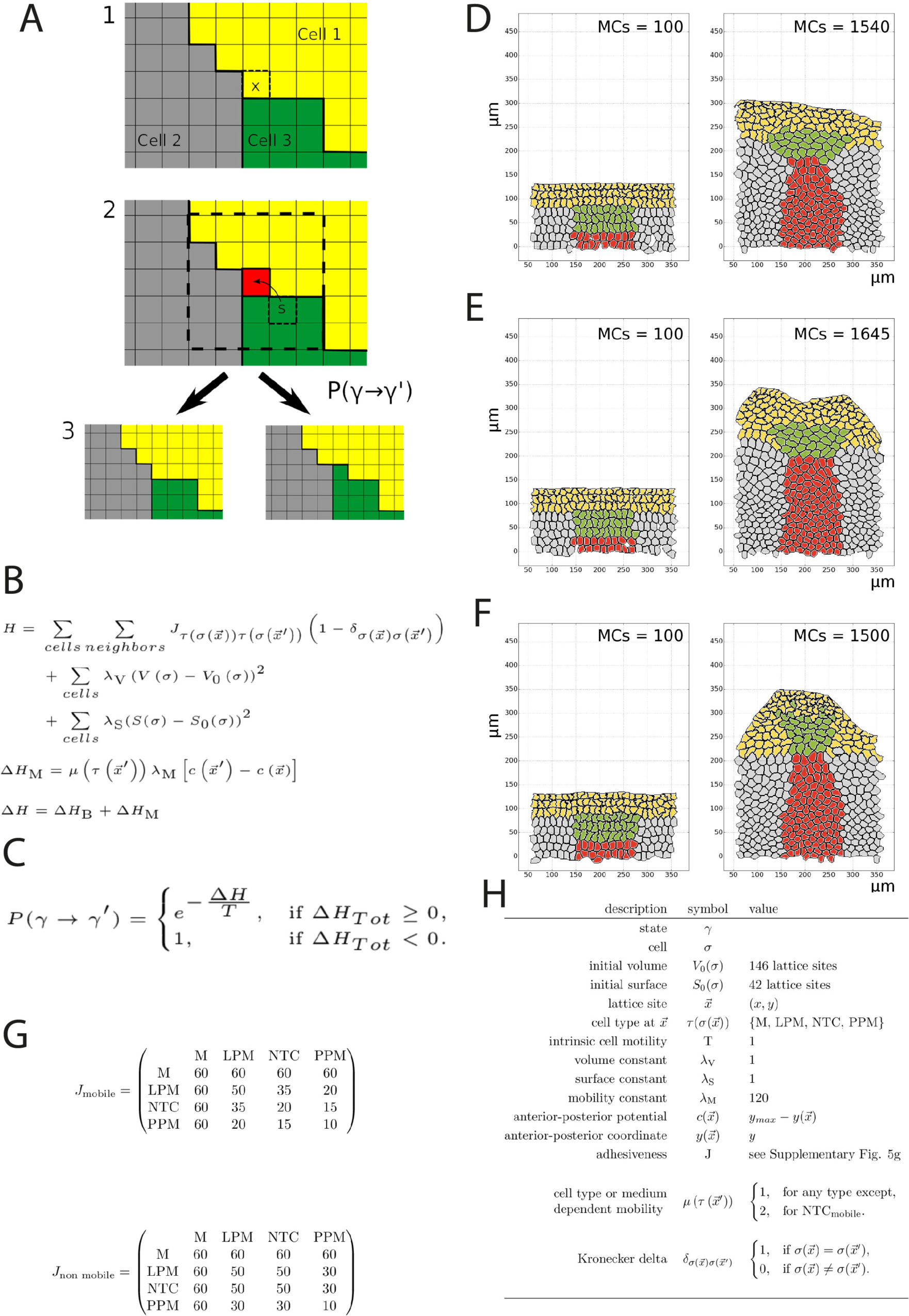
Cellular Potts model for tissue dynamics in the mesoderm. A, Schematic representation of the simulation protocol using three cells. (1) First, a random lattice site at the surface of a cell is chosen (x). (2) Next, either empty medium (M) or a random source (s) is chosen within 2 lattice sites (bounded area) of the initial target site (red). The Hamiltonian for the hypothetical new state is calculated and evaluated within the bounded area and the difference ΔH computed. (3) Depending on the probability P from state γ to state γ’ one of the two possible outcomes is chosen, either the source lattice site is copied into the target, or the target remains the same. B, Definition for the Hamiltonian H and modifications to ΔH for our CPM. Exact parameter values are found in supplementary figure 8H. (1) The Hamiltonian for state γ consists of three sums and defines the total energy of the system. The first sum is running over each cell σ at a lattice site xand its neighbouring lattice sites 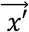. This sum represents the energetic contributions by the cellular adhesiveness J, which is dependent on the cellular types 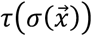 and 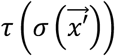. For the exact values of J, see supplementary figure 8G. The Kronecker-delta 5 makes sure that only different cells are contributing, and self-interactions are excluded. The second and third sum are running over each cell and sum up volume and surface contributions scaled by a factor λV, respectively λS. Each cell tries to preserve its original volume V0 and surface S0. (2) We defined a modification ΔHM to introduce mobility into our simulations with a linear anterior-posterior potential c. This modification is coupled to the mobility μ of the source cell at the neighbouring lattice site 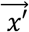 for the hypothetical state γ’ and the mobility constant λM. See supplementary figure 8A for a visual explanation of the simulation scheme. The total change in ΔH is obtained from both contributions of ΔHB = H(γ’) - H(γ) and ΔHM. c, The hypothetical state γ’ is accepted with a probability given by the Metropolis criterion. D-F, Simulation results, where every pixel represents one lattice site. M (white), PPM (yellow), NTC (green), LPM (gray), and NC (red). We indicate in the top right corner of each simulation the number of Monte Carlo steps (MCs). In each Monte Carlo step we loop through each surface pixel of every cell. The grid scale is 50µm. D, For the case of mobile leaders we see that no curving of the PPM takes place. The NTC take an oblong shape perpendicular to the direction of movement. The NC keeps a broad shape. E, In the case of adhesive leaders we see that the NTC still keep an oblong shape. A dip occurs in the PPM, resulting from the high attraction of the NTC. F, In the case of mobile & adhesive leaders we see a strong curvature of the PPM. The NTC and NC take a sharp and long configuration. G, values for adhesiveness as used in the simulation in matrix representation for the case of mobile leaders and non-mobile leaders. H, parameters as used in the simulation and for the definition of the Hamiltonian. We use a system in reduced units, where the cell motility is reduced to 1. The only free parameters are cell adhesiveness and mobility. The adhesiveness was chosen in such a way, that adhesive leaders stick roughly twice as strong together as non-adhesive leaders.

**supplementary figure 9.**
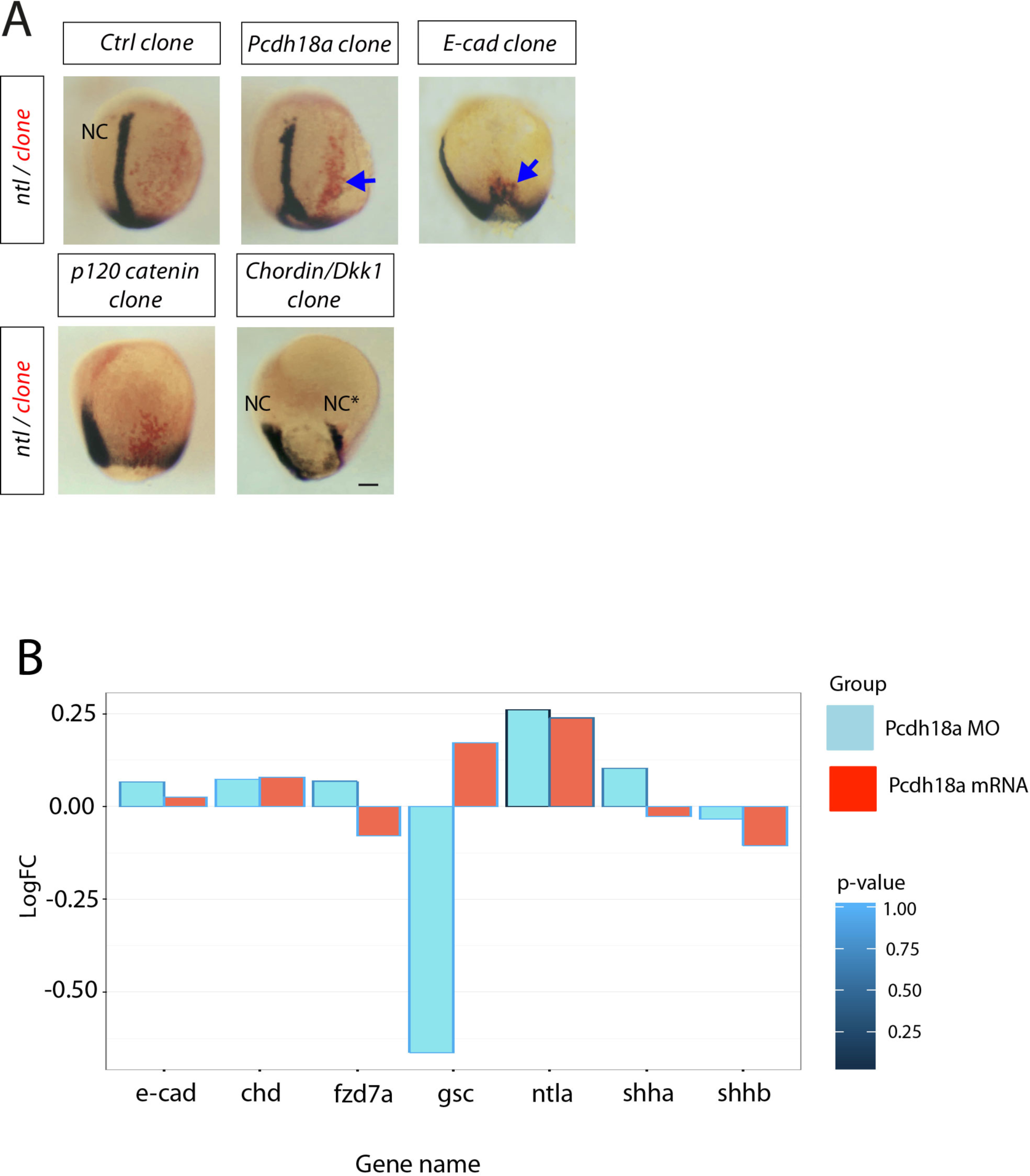
Pcdh18a induces a secondary notochord-like structure but lacks the transcriptional machinery of the shield organizer. A, Transplantation analysis of zebrafish embryos at 5hpf. Donor embryos were injected in the one-cell stage with indicated constructs and a lineage tracer. At 5hpf, a group of approx. 50 cells was transplanted into a WT host embryo in the lateral plate mesoderm. The embryos were allowed to develop to 9hpf, then fixed and subjected to ISH against ntl. Grafted cells were identified by the lineage tracer (red). NC, notochord; NC* ectopic notochord. Scale bar 100µm. B, Transcriptional profiling of zebrafish embryos injected with 0.1ng *pcdh18a* mRNA or with Pcdh18a MO at 24hpf. Limited alteration in the expression levels of genes known to influence Spemann organizer formation and function such as notochord markers (i.e. *gsc, goosecoid, shh-a, shh-b)*, or nodal/BMP/Wnt pathway related genes (i.e. *chd, chordin)* was observed.

